# Heterogeneity in ligand-bound TRPV1: A comparison of methods in cryo-EM and molecular dynamics simulation

**DOI:** 10.1101/2024.10.07.617120

**Authors:** Miro A. Astore, Robert Blackwell, David Silva-Sánchez, Pilar Cossio, Sonya M. Hanson

## Abstract

Cryogenic electron microscopy (cryo-EM) has emerged as a powerful method for resolving the atomistic details of cellular components. In recent years, several computational methods have been developed to study the heterogeneity of molecules in single-particle cryo-EM. In this study, we analyzed a publicly available single-particle dataset of TRPV1 using five of these methods: 3D Flexible Refinement, 3D Variability Analysis, cryoDRGN, ManifoldEM, and Bayesian ensemble reweighting. Beyond what we initially expected, we have found that this dataset contains significant heterogeneity— indicating that single particle datasets likely contain a rich spectrum of biologically relevant states. Further, we have found that different methods are best suited to studying different kinds of heterogeneity, with some methods being more sensitive to either compositional or conformational heterogeneity. We also apply a combination of Bayesian ensemble reweighting and molecular dynamics as supporting evidence for the presence of these rarer states within the sample. Finally, we developed a quantitative metric based on the analysis of the singular value decomposition and power spectra to compare the resulting volumes from each method. This work represents a detailed view of the variable outcomes of different heterogeneity methods used to analyze a single real dataset and presents a pathway to a deeper understanding of the biology of complex macromolecules like the TRPV1 ion channel.

## 1 Introduction

The field of structural biology has been dominated by a drive for high-resolution structures. Cryo-electron microscopy (cryo-EM) has emerged as the leading technique for determining these high-resolution structures for larger biomolecules [1, 2]. Recently, the ability of cryo-EM to capture not just a single structure but a distribution of states has been attracting growing interest. Access to this information hidden in existing cryo-EM data has implications for understanding biomolecular mechanisms and treating disease. Heterogeneity has been known to exist in vitrified samples since the inception of cryo-EM [3–5]. However, it is an open question whether continuous landscapes of conformational heterogeneity and the populations of their representative states can be reliably recovered from these samples. Recent analysis techniques take advantage of the latest advances in machine learning and statistical inference and range from geometric machine learning [6] to variational autoencoders [7–9] to flow field generators [10, 11] to linear or more classical Bayesian inference approaches [12– 14], we point the reader elsewhere for a more comprehensive review [5]. However, a rigorous comparison of the performance of these methods on a real dataset of biological interest is yet to be performed. Therefore, questions such as whether all methods reconstruct the same motions, how to quantitatively compare the results from different heterogeneity methods, and how they address compositional versus conformational heterogeneity remain to be addressed.

Here we have chosen a classic example in single particle cryo-EM — the tetrameric TRPV1 ion channel — to compare heterogeneity methods. As both the first molecular temperature sensor discovered in humans, and one of the first proteins solved to near-atomic resolution in single-particle cryo-EM, TRPV1 has historical significance to the field [15, 16]. The advancement of heterogeneity methods in cryo-EM holds great promise for further studies of this important molecular target, as heterogeneity is thought to play a significant role in the poorly understood mechanisms of the temperature-sensitive properties of the TRP family of ion channels [15, 17–21]. Unlike the temperature-sensing mechanism, the ligand-dependent activation of TRPV1 has been well-characterized [21]. For example, in a ligand-bound open state, when TRPV1 is bound to both double-knot toxin (DkTx) and resiniferatoxin (RTX), the cytosolic domains are shorter and wider than in the apo closed state. During the transition from an open to a closed state, the cytosolic domains and the transmembrane domains (TMDs) twist in opposite directions, and the cytosolic domains shift simultaneously downward, to change the height of the molecule. These concerted motions lead to a constriction of the pore, closing the ion channel (Figure 1C, Supplementary Movie 1). Despite the presence of high affinity ligands in the experimental dataset analyzed, a recent heterogeneity analysis method – 3D Flexible Refinement (3DFlex) – revealed deformations consistent with this transition from the ligand-bound open state to the apo closed state [10]. However, it is unclear whether the deformations suggested by explorations of the 3DFlex latent space correspond to real states in the sample, as the purpose of 3DFlex is to increase the resolution of the consensus refinement. This is a general problem with heterogeneity methods in cryo-EM. Usually these methods must embed particle images in a difficult to interpret high dimensional latent space, which can give rise to hallucinations. Experiments have recently been performed with synthetic data where heterogeneity was detected by several methods, even in a dataset with only a single molecular state [13].

**Fig. 1.**
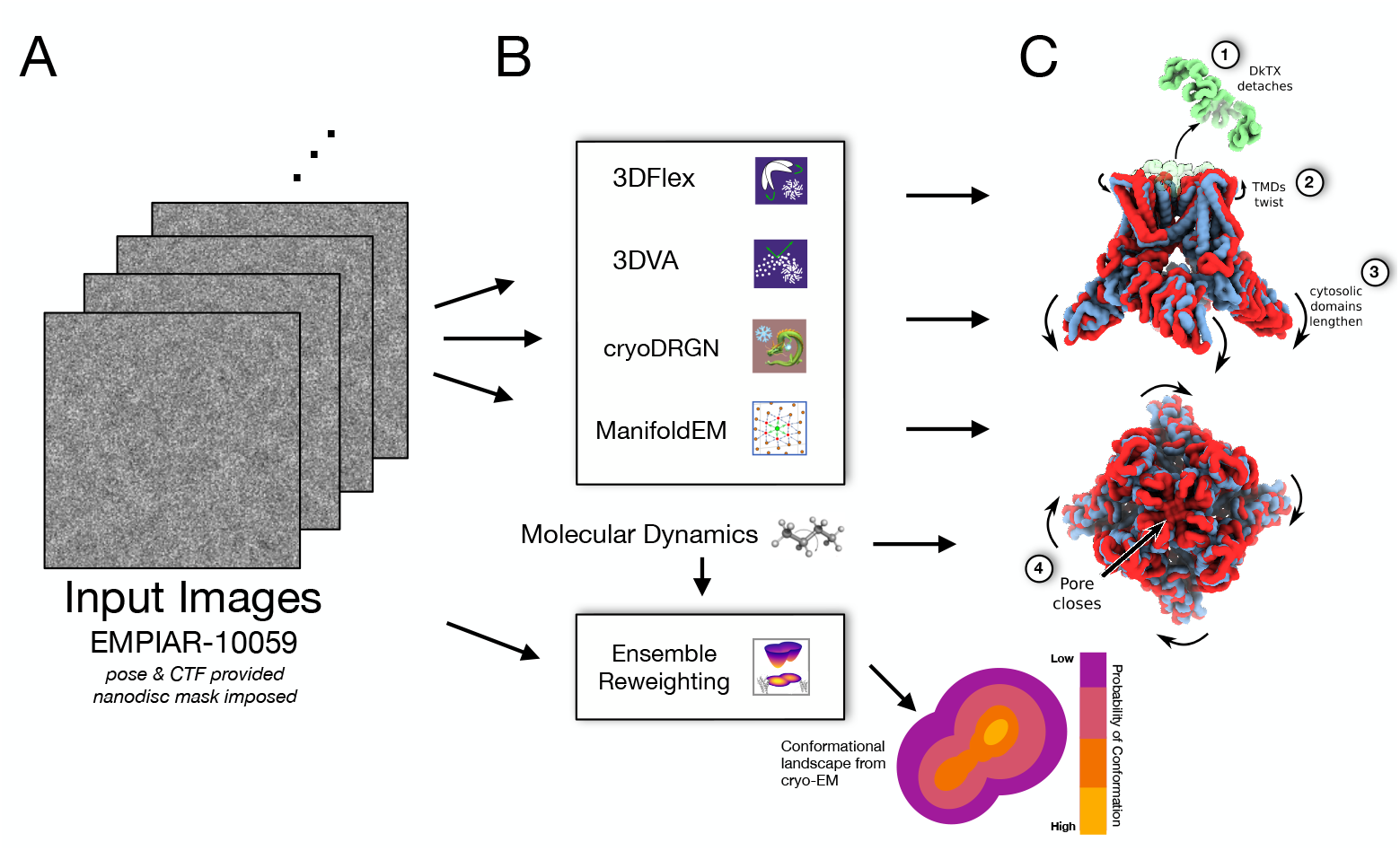
Overview of heterogeneity methods used to analyze the toxin-bound TRPV1 cryo-EM dataset found in EMPIAR-10059. A) For all methods the same set of 218,787 input images were used to determine if the same conformational and compositional changes were consistently picked up. Pose and CTF parameters were determined via 3D refinement in cryoSPARC and the nanodisc was masked out. B) Four methods for analyzing heterogeneity in cryo-EM and molecular dynamics simulations together with Bayesian ensemble reweighting were compared, each with its icon shown here. C) We have defined four types of conformational changes to look for: 1) presence or absence of the double-knot toxin (DkTx), 2) transmembrane domain (TMD) twist, 3) cytosolic domain lengthening, and 4) pore opening and closing. The molecular dynamics simulations were also compared to the cryo-EM data using Bayesian ensemble reweighting, which reweights the relative populations of atomic structures using the experimental data.

Thus, we were motivated to analyze this ligand-bound TRPV1 datset with the most well-established methods in conformational heterogeneity in single particle cryo-EM: 3DFlex [10], 3DVA [14], cryoDRGN [7], and ManifoldEM [6]. We found that each of these methods showed sensitivity to different aspects of heterogeneity in unexpected ways. For example 3DFlex proved sensitive to the detection of only conformational changes, while cryoDRGN is heavily favored toward compositional heterogeneity. Conversely, 3DVA and ManifoldEM were capable of detecting both kinds of heterogeneity. Despite these differences, each method consistently indicated the presence of a population of particles occupying states closer to the apo closed state, rather than the ligand-bound open state we initially expected to dominate the results. Furthermore, we ran atomistic molecular dynamics (MD) simulations of the ligand-bound TRPV1 system and found motions visually similar to those suggested by the cryo-EM analysis tools. Validating the results from disparate heterogeneity methods is an ongoing point of discussion within the field. However, with the development of a custom analysis pipeline based on singular value decomposition, we verified that the results from these disparate methods agree with one another—while we used the method of Bayesian ensemble reweighting to confirm the presence of these alternative states within the sample itself [12]. Overall, this case study of TRPV1 has consistently revealed heterogeneity in a dataset that would conventionally be expected to only contain a single state. For each method applied, we discovered many idiosyncrasies—each had a different sensitivity for either conformational or compositional heterogeneity. However, by using different methods in combination with each other, we were able to gain a more complete picture of what was present in the sample.

## 2 Results

### 2.1 3DFlex reveals TMD twisting and cytosolic domain lengthening

For all analyses in this study we used the 218, 805 particle images of TRPV1 bound to DkTx and RTX available in EMPIAR-10059 [20]. Following signal subtraction of the nanodisc (see Methods for details), we reconstructed this stack to a resolution of 3.0 Å using cryoSPARC. Recently, 3D flexible refinement (3DFlex) was added to the cryoSPARC software suite. It was designed to increase the resolution of flexible portions of a biomolecule by accounting for their conformational heterogeneity [10]. By embedding the molecule’s consensus reconstruction in a flexible mesh grid, 3DFlex introduces continuous deformations to a volume. The nature of these deformations are dictated by a generative neural network that uses a flow generator to convert a latent coordinate vector into a deformation field that convects the canonical map of the consensus reconstruction. It is trained to increase the overlap between a deformed volume and the electron density in particle images, thereby increasing the signal to noise ratio in flexible domains of the molecule. 3DFlex models are very sensitive to the choices of hyperparameters. For example, more rigid models exhibited noticeable asymmetries in their reconstruction of different subunits of the cytosolic domains, despite these domains being symmetric (Figure S1A). These asymmetries would persist on symmetry expanded particle stacks, and with models trained with a less rigid model (Rigidity = 1), but they were less pronounced. Further, we found that by independently training two models with the same hyperparameters, we could obtain noticeably different reconstructions and learned deformations (Figure S1B, Supplementary Movie 13). Despite these noticeable differences, these models yield the same training scores and global resolutions, raising questions surrounding the validity of 3DFlex-derived motions.

The original 3DFlex paper analyzed the TRPV1 dataset we focus on here, and showed increased local resolution of regions of the cytosolic domains that had previously been poorly resolved. In our hands, we achieved similar improvement in the local resolution of these cytosolic domains, to below 3 Å in some helices (Figure 2A-B). By training a two dimensional latent space, we find two dominant conformational changes in the sample (Figure 2C). The first is a deformation of the TMDs and cytosolic domains twisting in opposite directions, while the latter domains simultaneously lengthen, similar to what one would expect for a transition between the ligand-bound open and the apo closed states (Figure 2D, Supplementary Movie 2). The density increases by a height of approximately 5 Å. The second motion characterized by our 3DFlex model corresponds to an asymmetric squeezing motion of the cytosolic domains, with the helices moving approximately 5 Å (Figure 2E). Overall the motions we have seen are similar to those seen in the initial 3DFlex analysis of this dataset, though we found the optimization of hyperparameters nontrivial. Since the primary objective of this method is to increase the resolution of the consensus refinement, and because the extra density of the toxin may cause artifacts in the volume-preserving flow field, we were motivated to use other methods to investigate the existence of the motions seen in this ligand-bound TRPV1 dataset by the 3DFlex analysis.

**Fig. 2.**
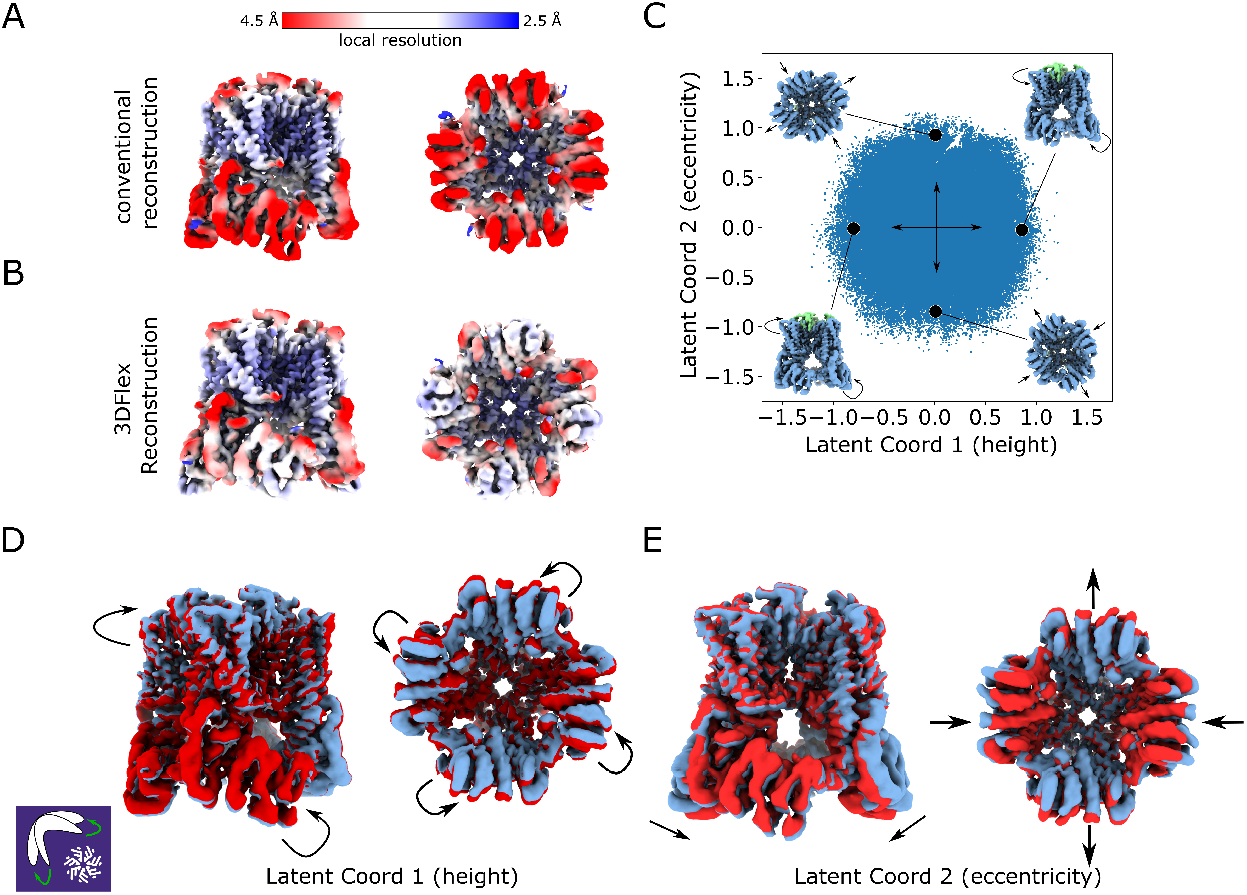
3D Flexible refinements accounts for flexible motions to improve their resolution. A) A local resolution estimation for conventional refinement of TRPV1. The cytosolic domains are poorly resolved. B) Local resolution estimation after 3DFlex refinement. Significantly more features in the cytosolic helices can be confidently resolved. C) Particles distributed throughout the latent space of the 3D Flexible refinement model, and visualizations of the deformations which these dimensions correspond to D) Visualization of the first deformation found by 3DFlex. The cytosolic domains and the TMDs twist in opposite directions, bringing the channel closer to either a closed or an open state. E) Visualization of the second mode found by 3DFlex. The cytosolic domains undergo an asymmetric squeezing motion.

### 2.2 3DVA detects TMD twisting, cytosolic domain lengthening, and toxin unbinding

3D Variability Analysis (3DVA) is a simpler analysis tool than 3DFlex, and is also available in cryoSPARC. It performs principal component analysis (PCA) on particle images in Fourier space [14], introducing deviations from the average volume along linear eigenvectors, to minimize the difference between volumes deformed along these eigenvectors and the ensemble of images. When applied to this TRPV1 dataset, 3DVA detects conformational changes comparable to those seen in 3DFlex: the first motion detected by 3DVA is the asymmetric squeezing of the cytosolic domains (Figure 3A), while the second motion is the twisting of the transmembrane and cytosolic domains (Figure 3B, Supplementary Movie 4). Note the different order compared to 3DFlex (Figure 3C). Additionally, the results from 3DVA appear to be robust to changes in hyperparameters. Changing the filter resolution of the 3DVA or choosing to expand the symmetry of the particle stack did not noticeably change our results. Adding larger numbers of latent dimensions did reveal small conformational changes, with higher amounts of asymmetry compared to the two studied in detail in Figure 3. These motions appear as a ‘wobbling’ of the cytosolic domains relative to the TMDs.

**Fig. 3.**
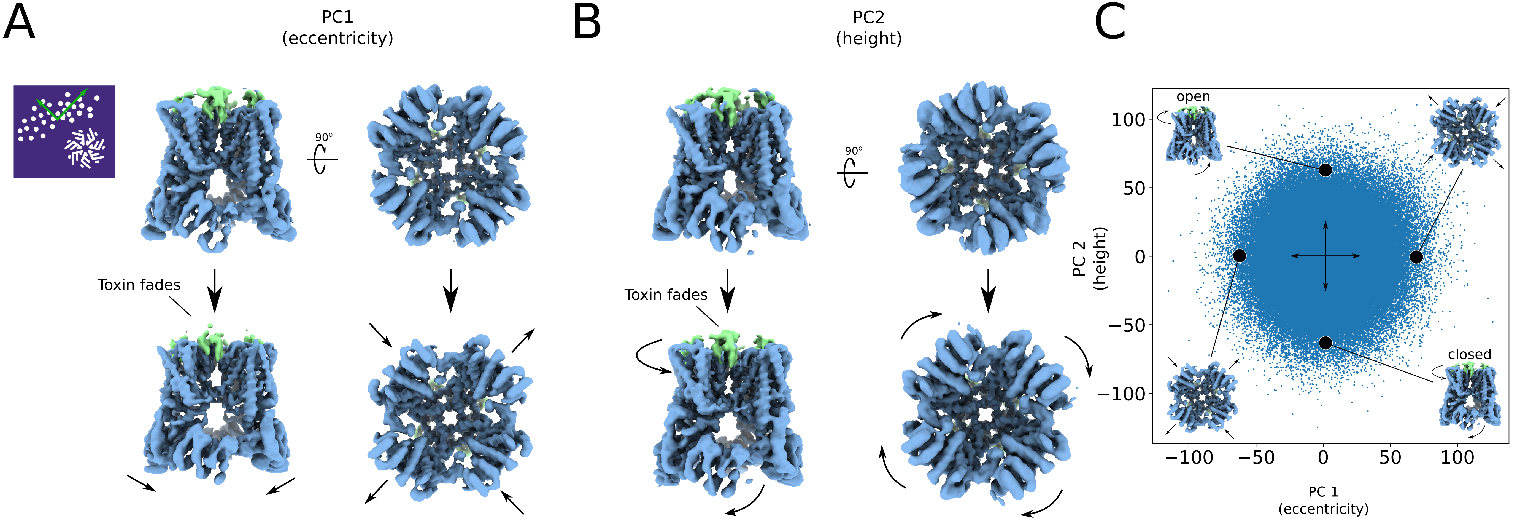
3D Variability analysis is sensitive to both compositional and conformational heterogeneity. A) Visualization of the first mode found by 3DVA. This mode captures asymmetric motions of the cytosolic domains and a change in occupancy of DkTx. B) The second mode found by 3DVA. This mode captures twists in the TMDs and changes to the height of cytosolic domains, bringing the channel closer to a closed state. There is also a change in the density of the DkTx toxin, but it is smaller than the change found in the first mode. C) Mapping the images into the 3DVA latent space indicates a Gaussian distribution.

In addition to the conformational changes detected by 3DVA, explorations of the latent space reveal evidence of compositional heterogeneity (Figure 3A-B). Both the first and second components of the latent space reveal changes in the density of the DkTx toxin, indicating that there is compositional heterogeneity in this particle stack—in some particles, the toxin has partially or fully unbound from the protein. Given its fast runtime (Supplementary Table 2) and reliable results, we found that 3DVA could be used to quickly identify the main sources of heterogeneity in a dataset. However, the basis of linear vectors in a Fourier basis may influence which kinds of molecular changes are detected [22].

**Table 1.** Summary of conformational and compositional heterogeneity detected by each method.

| Method \ Conformational change | Pore closing | TMD twist | CTD Motions | Toxin Occupancy |
| --- | --- | --- | --- | --- |
| 3DFlex | × | ✓ | ✓ | × |
| 3DVA | × | ✓ | ✓ | ✓ |
| ManifoldEM | ✓ | ✓ | ✓ | ✓ |
| CryoDRGN | × | × | × | ✓ |
| MD Simulations | ✓ | ✓ | ✓ | × |

**Table 2.** Computational cost of heterogeneity methods. * = computations performed on a single A100 GPU. ** = computations performed on four A100 GPUs in parralel. *** = computations performed on a 16-core Xeon Gold 6234 CPU@3.3 GHz.

| Method | Computational wall time |
| --- | --- |
| 3DVA | 25 minutes * |
| 3DFlex | 4 hours * |
| CryoDRGN | 35 hours (20 minutes per epoch) ** |
| ManifoldEM | 6 hours *** |

**Table 3.** Summary of all analysis performed on EMPIAR-10059.

| Analysis Tool | Nanodisc mask | Symmetry Expansion | Picking Extra Particles | Filtered particles [33] |
| --- | --- | --- | --- | --- |
| 3DFlex | ✓ | ✓ | ✓ |  |
| 3DVA | ✓ | ✓ | ✓ | ✓ |
| CryoDRGN | ✓ | ✓ | ✓ | ✓ |
| Manifold Embedding | ✓ | ✓ | ✓ |  |

### 2.3 CryoDRGN is more sensitive to compositional heterogeneity than other methods

While 3DFlex and 3DVA have the convenience of being implemented in cryoSPARC another popular tool for analyzing conformational heterogeneity in single particle cryo-EM is cryoDRGN (Deep Reconstructing Generative Networks) [7]. CryoDRGN is a variational autoencoder (VAE) that consists of two neural networks built with an image-encoder-volume-decoder architecture. The first network takes raw particle images as input, and outputs a low dimensional vector, embedding the images in a latent space. The second neural network takes a point in this latent space as input and outputs a 3D volume. These neural networks are trained such that reconstructions from images in a specific region of this latent space will produce densities that are similar to the volumes predicted by the decoder network.

Applying cryoDRGN to this TRPV1 dataset resulted in a latent space that captured significant changes to the occupancy of DkTx, similar to what was seen in 3DVA (Figure 4A, Supplementary Movie 6). However, there are comparatively few conformational changes to the TRPV1 protein itself, unlike 3DVA and 3DFlex. By identifying regions of the latent space which produce volumes with little DkTx bound, we can identify clusters of particles which are likely to have a low presence of DkTx (Figure 4B). Direct reconstructions of images in these clusters revealed volumes with toxin occupancy consistent with the predictions of the cryoDRGN VAE (Figure 4C, Supplementary Movie 9). This indicates that some particles in the sample do not have DkTx bound to TRPV1, or that some particles have DkTx bound to only certain subunits. Given our other results the lack of conformational heterogeneity in the cryoDRGN result was unexpected, and suggests that compositional heterogeneity can overwhelm small conformational changes with this VAE architecture.

**Fig. 4.**
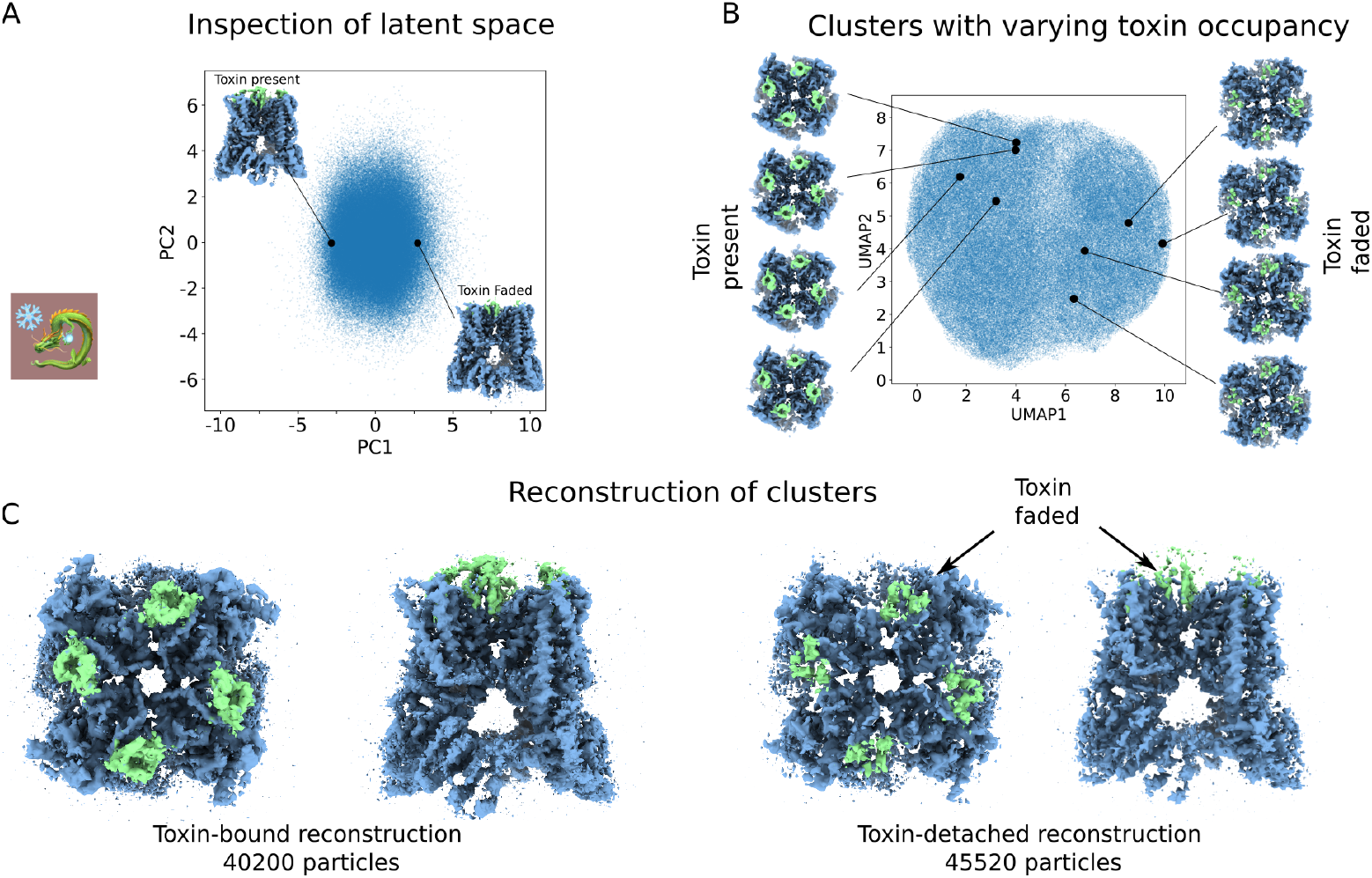
CryoDRGN is sensitive to compositional heterogeneity. A) The most variation in the cryoDRGN latent space occurs in the variation of the occupancy of the DkTx toxin, bound to the extra cellular face of TRPV1, with relatively little conformational change in the protein itself. B) Clusters in the latent space of the cryoDRGN model suggest toxin occupation at different levels, giving rise to compositional heterogeneity. C) Selection of particles surrounding the parts of the latent space around these clusters yields reconstructions consistent with the visualizations of the cryoDRGN decoder network.

### 2.4 ManifoldEM resolves small motions

Manifold embedding, or ManifoldEM, represents one of the first approaches to study of conformational heterogeneity in cryo-EM [6, 23]. This method has been used to resolve conformational transitions in the SARS-COV2 spike protein [24, 25], the ribosome [6], and the ryanodine receptor [26]. However, despite previous versions of the code being made available [26–28], the methods adoption was not wide spread due to its slow speed and lack of stability. In this work we have used a modern Python version of ManifoldEM which is two orders of magnitude faster and actively maintained (see Methods for details). Unlike other methods, ManifoldEM first analyzes images from the same projection direction (PrD) before stitching the results together over the full orientation sphere. After an initial Euclidean distance calculation followed by diffusion mapping, non-linear Laplacian spectral analysis (NLSA) is then performed to project the diffusion map results back into the interpretable space of 2D images. Initial versions of ManifoldEM then required the user to manually annotate the resulting 2D eigenvectors for every PrD, but a more recent development allows this process to be semi-automated with only a few ‘anchor nodes’ defined by the user and belief propagation of the motions determined by the correspondence of optical flow vectors [28]. Together, these algorithms determine the global conformational coordinate for the motion of interest in three dimensions.

The focus on projection directions allows ManifoldEM to detect motions not visible to other methods. In top views of TRPV1, we could detect changes to the size of the pore of the ion channel (Figure 5A). While in side views, motions in both the cytosolic domains and transmembrane helices were visible (Figure 5B). These latter results are consistent with the descriptions from the conformational changes suggested by other methods. While these motions appear in the ManifoldEM analysis per PrD, the volumes produced in the final stages of the analysis are not high enough resolution to capture them, likely because any PrD with fewer than 200 images are not included, resulting in roughly half of the total particle stack to be discarded. Even though they are lower resolution than other methods, the trajectory of volumes at different locations of the ManifoldEM latent space display changes in the occupancy of the DkTx toxin, similar to the results of cryoDRGN (Figure 5C-D, Supplementary Movie 11). Intriguingly, these compositional changes were not obvious in the individual NLSA eigenvectors of individual PrDs.

**Fig. 5.**
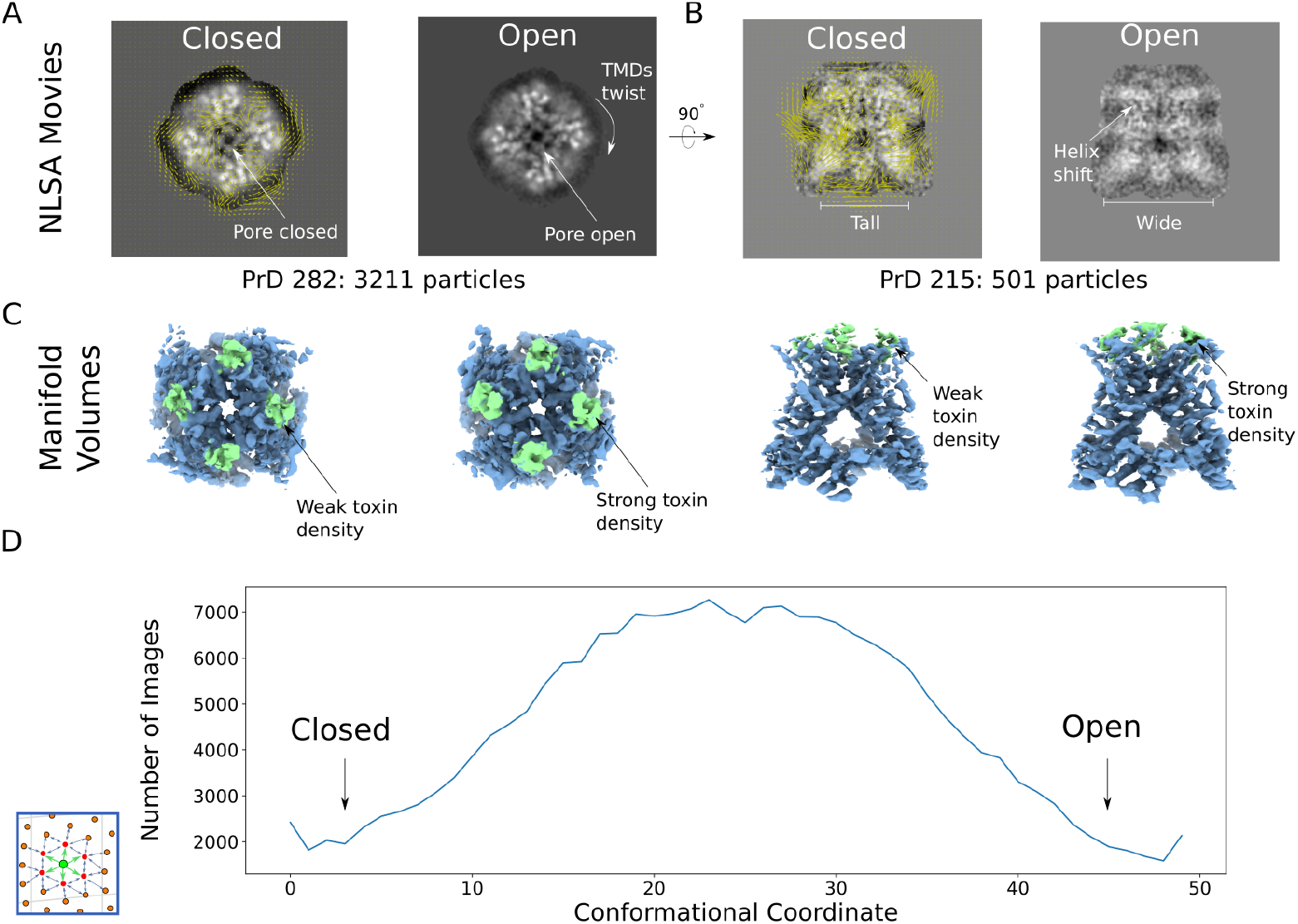
ManifoldEM resolves small motions. A) A top down view of the optical flow calculated of a representative PrD from the ManifoldEM results. This view reveals twists in the TMDs and the pore in both an open and closed configuration. B) A side-on view of the optical flow calculated of representative PrD from the ManifoldEM result. This view reveals heterogeneity in the nanodisc and motions in the cytosolic domains. C) The distribution of states along the conformational coordinate. The distribution closely resembles a Gaussian, similar to the latent spaces discovered by cryoDRGN, 3DFlex and cryoDRGN.

### 2.5 MD simulations reveal motions similar to those found in cryo-EM

The methods we have studied in previous sections have relied solely on the analysis of cryo-EM images. They made no assumptions about the underlying physics of molecule under study. By contrast, numerical simulation techniques like molecular dynamics use physical priors to generate a distribution of states for a biomolecule. Hence, we ran atomistic molecular dynamics (MD) simulations of the TRPV1 homotetramer bound to DkTx and RTX in a POPC bilayer as an orthogonal method to compare the motions found by the previsely-described image-based methods (Supplementary Figure S2). Analyzing 5×2*μs* simulations run at 310 K, the eigenvectors discovered by applying a version of principal component analysis (PCA) adapted for the analysis of heterogeneous symmetric particles reveal deformations to the protein to those seen with heterogeneity analysis of the cryo-EM images (Supplementary Movie 12). In the MD simulations, however we have more precise access to the atomistic detail of these asymmetric motions and their dynamics over time, as can be seen in the plots of these motions in individual trajectories (Figure 6). Overall, even with the toxins bound throughout the MD simulations, we see the TMD twist and domain motions that indicate an oscillation between the existing toxin-bound open and apo closed state structures originally resolved from the cryo-EM structures (Supplementary Movie 12). We summarize this and the results of the cryo-EM heterogeneity methods in (Table 1). This table demonstrates the correspondence between our physics-based MD simulations and the image-based cryo-EM methods.

**Fig. 6.**
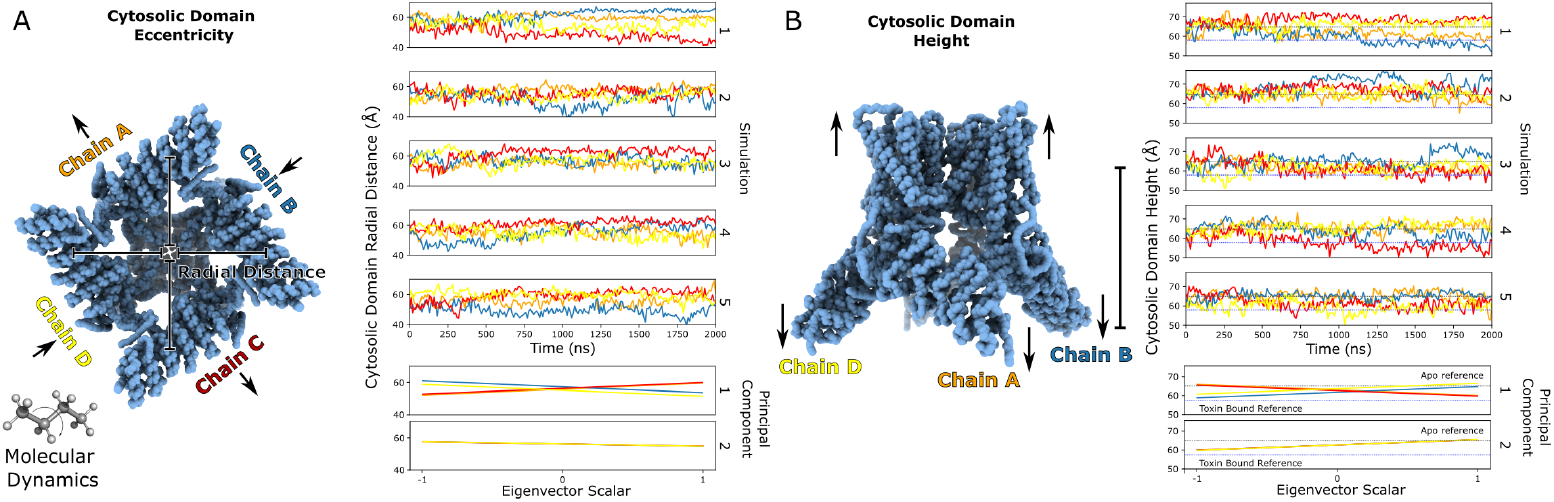
MD reveals motions similar to those found in heterogeneity analyses of the cryo-EM images. A) Changes in the distance of the cytosolic domains to the center of protein, colored by chain. These changes to the eccentricity of the molecule are best captured by the first principal component of the molecular dynamics trajectory. The DkTX toxin has been removed from all analysis of the simulations. B) Changes to the overall height of TRPV1 where the TMDs, colored by chain. These changes are best captured by the second principal component of the molecular dynamics trajectory which also includes a twisting of the TMDs.

### 2.6 Ensemble reweighting shows a distribution of structures in the cryo-EM data

So far we have been comparing the results of heterogeneity analysis methods and MD simulations by visual inspection of the conformational changes of TRPV1. However, with the MD simulations now in hand, we have an avenue for a more a quantitative approach to comparing the MD and cryo-EM images—Bayesian ensemble reweighting. In this method, the likelihood that a given structure came from a given image is calculated [12, 29]. The prior estimation for the populations of different structures, which comes from MD simulations, is reweighted to reflect the populations of this structural ensemble in a given set of cryo-EM images. In this case we use the PCs from MD, as described above, as our state space. Indeed, reweighting the MD distribution to that of the cryo-EM data indicates that the majority of particles occupy a conformation close to the ligand-bound open state, but with a small population of particles closer to the apo closed state (Figure 7A, B).

**Fig. 7.**
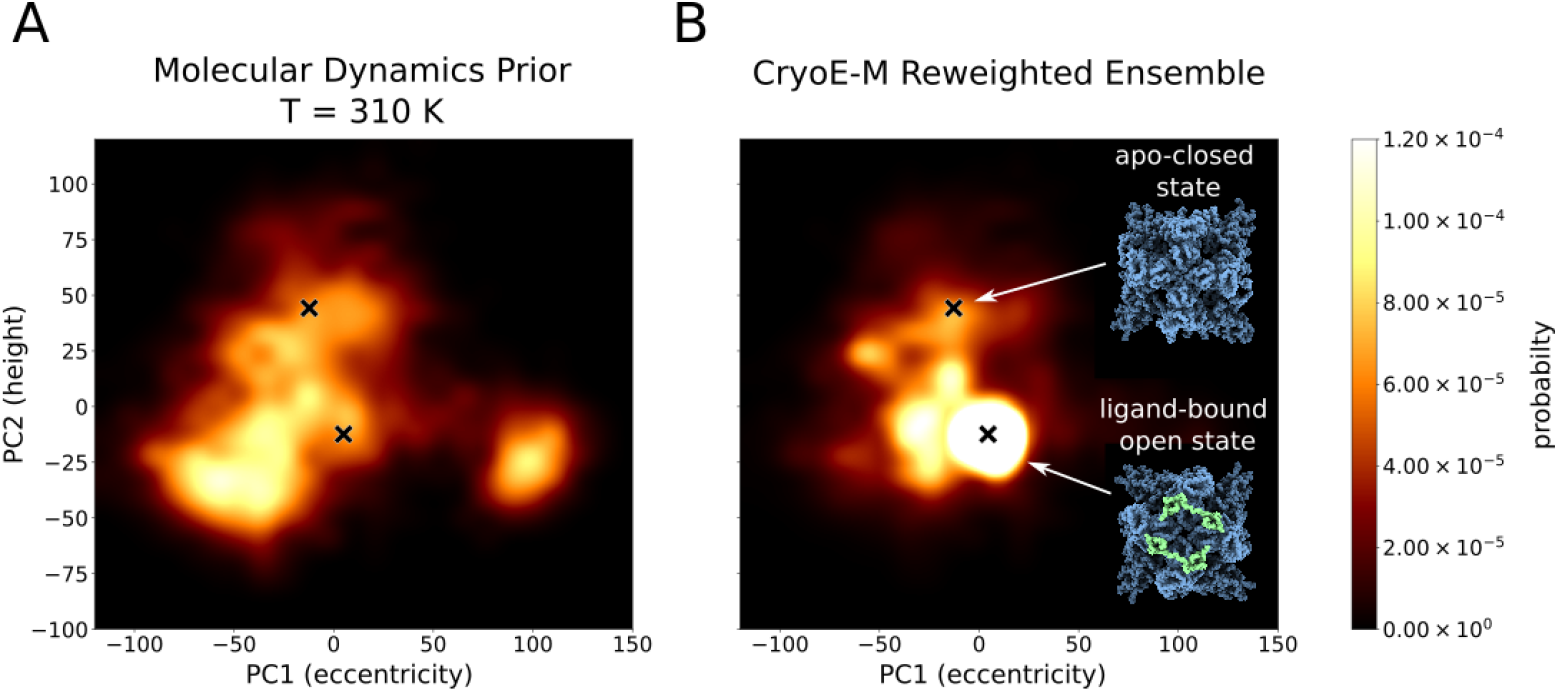
Ensemble reweighting confirms the presence of structures closer to the closed state. A) The MD density landscape at 310K along the principal components. B) The cryo-EM reweighted landscape using the particle images and structures from molecular dynamics simulations. The locations of the apo-closed state and ligand-bound open state are indicated. No DkTX toxin was bound to the models of the TRPV1 protein during the analysis shown here. The results reflect the internal conformational changes of the protein.

One advantage of ensemble reweighting is that it represents the distribution directly in the interpretable conformation space defined by the MD simulations. This is in contrast to the latent spaces of other heterogeneity methods which resemble simple Gaussian distributions. The reweighting method allows us to access higher resolution features at will—something that other methods, which derive their latent spaces from images alone, will struggle to do [13].

Another remarkable feature of the ensemble reweighting is that the distributions from ensemble reweighting seem to converge with a relatively small number of particles: only 5% of the total number of particle is necessary to find which states are present and with only 23% of the total number of particles, the weights of the different structures have converged (Figure S3A). One nuanced aspect of the ensemble reweighting method in cryo-EM is the choice of which specific structures to use to represent the full MD ensemble. Indeed, we notice that when structures from all three temperatures at which we ran simulations (150 K, 200 K, and 310 K) are used in reweighting, the structures derived from lower temperature simulations received higher weights (Supplementary Figure S3B-C, Supplementary Figure S4). Indicating that the sample ensemble may be significantly cooler than the temperature the sample’s initial temperature 277 K (4°). These results show that ensemble reweighting is able to detect rare states in this sample, despite the fact that the simulations are carried out using classical forcefields at different temperatures from the incubated temperature of the sample using.

### 2.7 Comparing heterogeneity methods with Singular Value Decomposition and spectral analysis in CryoSVD

Ensemble reweighting allows us to compare the MD simulations to cryo-EM images directly. However, it is also desirable to compare the volumes produced by different heterogeneity methods with each other. We have designed a pipeline for this purpose based on simple linear algebra which we call CryoSVD. We first select one or two volume series per method *e*.*g*. a trajectory of volumes along PC1 for 3DVA. Each volume is then masked, normalized, and the mean volume is subtracted from each member of the series. Singular Value Decomposition (SVD) is then performed on the series of processed volumes, resulting in a basis for each individual series. Then, we take these eigenvectors and perform a second round of SVD to find a common basis for all the series. By doing two rounds of SVD, we can compare the different series both quantitatively and qualitatively (Figure 8A). The first round lets us compare series quantitatively. When we project the eigenvectors from each series onto one another, we can calculate a quantity known as Proportion of Captured Variance (PCV) [13]. This measures how similar two given basis sets are, with a PCV of 1 indicating that the basis sets are identical, while a PCV of 0 indicates that all the basis vectors in the two sets are orthogonal. The second round of SVD allows us to compare the series qualitatively. By projecting volumes into the common basis, we can see whether they map to similar regions in the shared space.

**Fig. 8.**
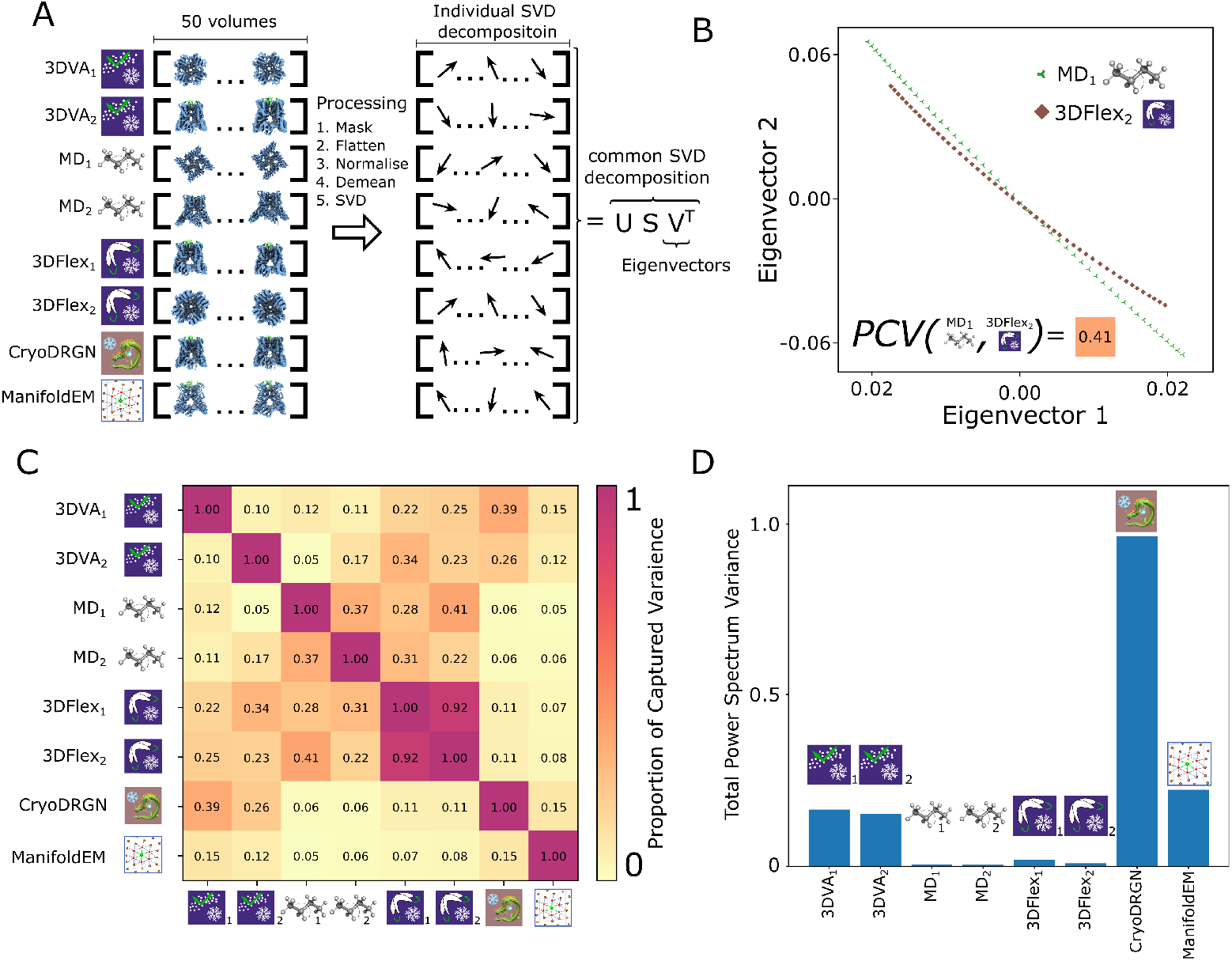
Comparing series of volumes in a linear basis. A) To compare series of volumes, we normalize them and perform singular value decomposition on each series individually. These basis vectors obtained from this process are then used to calculate a common basis in another round of SVD. B) The coefficients of a given series in this common basis allow us to compare the paths traced by multiple volume series. C) By projecting basis of one volume series onto another, we can quantify the similarity between the series in a quantity known as proportion of captured variance (PCV). The closer this quantity is to 1, the more similar the volume series are. D) Inspections of the power spectra for volume series as a means to quantify the amount of compositional heterogeneity captured in the series. When there is compositional heterogeneity a volume series, there will be more variance in the power spectra of its constituent volumes.

Using this pipeline, we can clearly see that volumes sampled along 3DFlex latent 2 are closely related to volumes generated from PC1 of our MD simulations. They form remarkably similar trends when projected onto the first two eigenvectors of the common basis, and their PCV is 0.41. This is the closest match we have found between two series from different methods (Figure 8B). This is consistent with our observations in section 2.5, confirming that both these volume series capture an asymmetric squeezing of the cytosolic domains of TRPV1.

A pairwise comparison of all methods using PCV demonstrates that series which capture the same qualitative changes to the molecule have more similar SVD bases (Figure 8C). In general, series which capture the asymmetric squeezing of the cytosolic domains share higher PCV scores with each other (MD mode 1, such as 3DVA mode 1, and 3DFlex mode 2) than with the series which capture changes to the height of the molecule (MD mode 2, 3DVA mode 2 and 3DFlex mode 1). Similarly, we see that series which capture changes to the occupancy of the DkTX toxin (cryoDRGN, ManifoldEM, and 3DVA) have higher PCV scores with each other than with series which do not capture changes in the toxin (MD simulations and 3DFlex). These trends are also apparent from inspecting the path of all series in the common basis (Supplementary Figure S8).

With the SVD pipeline demonstrated here, it can be difficult to distinguish between conformational and compositional heterogeneity. To study compositional heterogeneity, one could filter the refined map to detect contrast loss, but it is not simple to expand this approach to a series of volumes [30]. For our purposes, we can inspect the radially averaged power spectrums of a volume series. Since translational degrees of freedom will be degenerate in the Fourier basis, a series which only captures conformational heterogeneity will exhibit very similar power spectrums in each volume (Figure 8D). Indeed, we can see that the methods which capture compositional heterogeneity from the DkTX toxin (ManifoldEM, cryoDRGN and 3DVA) have significant variation in their power spectra compared to methods which only capture conformational heterogeneity (MD simulations and 3DFlex).

## 3 Discussion and conclusions

Here we have presented our analysis of a canonical dataset of the TRPV1 ion channel using modern heterogeneity analysis tools. After the variety methods we have now applied to tease out the heterogeneity of this TRPV1 dataset, we believe the results shown by the 3DFlex team of the conformational heterogeneity of TRPV1 to be true motions exhibited by the molecule in both MD simulations and the cryo-EM data. This work has implications for the analysis of heterogeneity in cryo-EM for which a comprehensive comparison of methods on a real dataset has not been performed until now. While we believe other datasets with fewer complicating factors, such as compositional heterogeneity and nanodisc heterogeneity, are better suited to benchmarking and methods development [31, 32], the results presented here serve as a solid introduction to the ways in which resolving heterogeneity from single-particle cryo-EM datasets can be complicated by a variety of factors, such that the choice of method used to study a dataset will heavily influence the kinds of heterogeneity a user will resolve.

A major concern when analyzing datasets with heterogeneity is the bias induced by using a subset of particles from which a high resolution reconstruction can be obtained. Currently, this is a limitation of most heterogeneity analysis methods in cryo-EM as jointly estimating Euler angles and conformational variability remains challenging. While the EMPIAR entry for this dataset provides 218,787 extracted particles, only 73,929 were used for the final 2.95 Å resolution reconstruction [20]. The EMPIAR entry does not provide this subset of particles, but we performed our own 3D classification and found that even using the 57, 602 that result in a 3.04 Å reconstruction in our hands, 3DVA analysis shows similar heterogeneity to the full particle stack used in the analyses described here (Supplementary Movie 14). This is consistent with experiments we conducted using cryoSieve [33], with which we produced a particle stack of only 27,572 particles (see Methods for more detail: this stack was obtained after 8 iterations of the cryoSieve algorithm and reconstructs to a resolution of 3.6 Å with cryoSPARC) – and still we were able to resolve heterogeneity with 3DVA. Furthermore, we were curious if even the 218,787 particles in EMPIAR had already discarded some heterogeneous particles, and therefore returned to the micrographs to repick particles, obtaining a particle stack of 525, 910 images with increased our reconstructed resolution to 2.8 Å. However, we could not resolve any additional heterogeneity from this enlarged particle stack, with one exception – in the case of cryoDRGN, small motions emerged in the helices in the cytosolic domains that were not previously visible (Supplementary Movie 7). Overall these experiments with particle stack size lead us to believe that while large numbers of particles can help resolve heterogeneity more clearly, if there are heterogeneous states in a sample they can be detected with even a small subset of high quality images of the molecule in question. This is supported by the detection of numerous states in the sample from Bayesian ensemble reweighting (Supplementary Figure S3).

One of the motivations to analyze the heterogeneity in this cryo-EM dataset of TRPV1 is the potential biological relevance of the conformational heterogeneity of this temperature sensor, for which the molecular mechanisms of temperature sensitivity is still debated. Indeed that the cytosolic motions we have established here exist within the toxin-bound state, as seen in both MD simulations and the cryo-EM data, has implications for the relevance of mutants seen in the cytosolic domain that have drastic effects on the temperature sensitivity of the channel [34, 35]. The temperature sensitive motion of the cytosolic domain has been hinted at by previous MD simulations, but this study represents the first time these motions have been confirmed in the context of structural biology experiments [36–38].

One interpretation of the results described here is that TRPV1 seems to display significant conformational entropy around an average state, and that with traditional cryo-EM analysis methods only this average state can be resolved to high resolution. By interrogating datasets to understand the full conformational variability we can access more than this average state, especially relevant for any mutants that alter this conformational entropy that can be understood from the perspective of “dynamic allostery” [39, 40]. Furthermore, that we see some toxin unbound state in the cryo-EM data, as picked up by the cryoDRGN analysis, which seems biased towards picking up compositional heterogeneity, lends some ambiguity to the exact states present in the cryo-EM data. This is especially interesting, as the toxin concentration in the experiment exceeds saturating concentration, but we find that the toxin-bound particles have a non-uniform distribution of viewing angles as compared to the unbound particles (Supplementary Figure S7), indicating that perhaps the air-water interface is playing a role here.

With the access to conformational heterogeneity that we demonstrate here, it is tempting to start talking about extracting thermodynamic information from these datasets, such as free energy landscapes or even entropy as alluded to above. However claiming to have access to this information must be done with caution for a variety of reasons. One initial caveat is that the latent spaces output from these heterogeneity methods, may be divorced from the appropriate molecular coordinates that would allow for an appropriate free energy landscape of these molecules. This can sometimes lead to hallucinations in the results of some heterogeneity methods [13]. Furthermore, the distribution of the conformational space would only represent the free energy of the molecule within the thin film environment of the grid after vitrification. The effect of the vitrification process on the biomolecular ensemble represented in single-particle cryo-EM datasets is still unknown. One computational approach found that barriers higher than 10 kJ/mol should preserve the distinction between states during the vitrification process [41]. However, this result was derived for an optimistically fast cooling rate of the ensemble [42]. Finally, while methods for extracting heterogeneity from single-particle samples have been improving in recent years, it remains unclear how to interpret the density of states in their respective latent spaces. This is critical for using them to understand the thermodynamics of macromolecules [31, 32]. There is a crucial need for validation methods for the reconstruction of multiple states in a sample. In the case of the reconstruction of single structures, “resolution” calculated from half-sets serves this purpose [43, 44]. However, there is no accepted analogous metric for heterogeneous ensembles.

Overall, this study of a real TRPV1 dataset serves as a proof of concept for more ambitious experiments that can be carried out by exploiting the natural heterogeneity of vitrified samples in cryo-EM. It is also the first demonstration of Bayesian ensemble reweighting on a real dataset, without the need to choose a path collective-variable [12, 29]. All of these methods for capturing heterogeneity in cryo-EM, though with special emphasis on Bayesian ensemble reweighting, have significant promise beyond standard single particle cryo-EM, namely time-resolved cryo-EM [45, 46] and *in situ* cryo-EM and cryo-ET [47–49], where fewer particles and less control of the environment of the sample will necessitate better analysis techniques. Underlying all of these promising applications is the need for more rigorous validation metrics. The metrics we have derived here help us validate whether the results from different methods agree, but they do not quantify how confident we can be that the results reflect what is present in the sample. Improvements in this space will help push these techniques toward mainstream acceptance by the biophysics community for the characterization of biological macromolecules.

## 4 Methods

### 4.1 Preprocessing the single-particle cryo-EM image stack for heterogeneity analysis

#### 4.1.1 Signal subtraction of the nanodisc

For this study we analyzed the publicly deposited raw electron microscopy data at EMPIAR-10059 [20]. This dataset consists of micrographs and particle stacks of a nanodisc-embedded TRPV1 protein bound to two agonists: a peptide toxin known as double knot tarantula toxin (DkTx) and a vanilloid compound known as Resiniferatoxin (RTX). Since TPRV1 is a membrane protein, it was was reconstituted into lipid nanodiscs with MSP2N2 scaffolds prior to imaging. This forms nanodiscs with a diameter of roughly 150 Å, significantly larger than TRPV1 itself. Nanodiscs in general can form at different sizes in solution, leading to compositional heterogeneity between particle images of the nanodisc itself [50]. Additionally, it is thought that variations in the location of the protein within the nanodisc also results in apparent nanodisc heterogeneity. Consistent with these aspects of membrane protein reconstitution into nanodiscs, in several cases our initial heterogeneity analyses were dominated by the compositional heterogeneity of the nanodisc (Supplementary Movies 8, 3, 5). By creating a significantly dilated mask to account for the particles with larger nanodiscs, we were able to use signal subtraction to decrease the artifacts from this nanodisc heterogeneity. Of the algorithms we tested, cryoDRGN and 3DVA appeared to be the most sensitive to nanodisc heterogeneity (Supplementary Movies 5, 3). Similar to this trend, we noticed in ManifoldEM that signal subtraction and masking of images increased the amount of motion detected by the algorithm (Supplementary Movie 15). These results reflect the importance of careful preprocessing for the analysis of heterogeneity in single particle cryo-EM.

#### 4.1.2 Homogeneous reconstruction and particle picking

For all main figures, the source particle stack was the polished particle stack 218, 787 particles deposited in EMPIAR-10059. Using homogeneous and local refinement with C4 symmetry, we were able to reconstruct this signal subtracted particle stack to a resolution of 3.0 Å. However, experiments were also performed with an enlarged particle stack. To expand the count of particles, we referred back to the original micrographs in the EMPIAR deposition and utilized a blob picker with a radius between 200 and 300 Å. This yielded 598, 273 particles. With 2D classification, we discarded particles which were obviously junk, but kept particles in classes which would usually be discarded because of their low quality. This yielded 525, 910 particles in total. This enlarged stack reconstructed to a resolution of 2.82 Å using homogeneous refinement followed local refinement under C4 symmetry constraints.

We also performed experiments using drastically reduced particle counts using cryoSeive [33]. Our starting particle stack for this algorithm was the expanded particle stack mentioned above, and the consensus model from this particle stack was used as a reference model for the cryoSieve algorithm. Frequency marching began from 40 Å and 8 iterations of the cryoSieve algorithm were applied with a retention ratio of 80% between iterations. This resulted in a particle stack with 27, 572 particles, reconstruction resolution of 3.6 Å using C4 symmetry constraints with homogeneous and local refinement in cryoSPARC.

### 4.2 Heterogeneity Methods

#### 3D Flexible Refinement

The signal subtracted particles were down sampled to 128 pixels, resulting in a pixel size of 1.83 Å. A tetrahedral mesh of 754 points and 2517 cells was generated to cover the down sampled volume. The 3Dflex model itself consisted of 2 latent dimensions with 64 hidden units in the multiperceptron architecture. A rigidity (*λ*) of 1 and latent centering strength of 5 were applied to the model during training. No consideration of symmetry was given to the 3DFlex model during training unless otherwise specified. In order to produce a series of volumes for section 2.7, we sampled 50 volumes along each latent coordinate defined by the 3DFlex model in Figure 2.

#### 3D Variability Analysis

The particles were analysed using 3D Variability analysis after they were prepared using signal subtraction and local refinement. A filter resolution of 5 Å was used during all 3DVA calculations. Unless otherwise stated, no consideration of symmetry was given for 3DVA calculations. In order to produce a series of volumes for section 2.7, we sampled 50 locations along the first and second latent coordinates of the 3DVA model defined in Figure 3.

#### CryoDRGN

CryoDRGN was trained with the default parameters. For the underlying VAE a latent space of 8 dimensions was used. Both the encoder and decoder networks consisted of 3 hidden layers with 1024 nodes each. The activation function for the VAE was the ReLU function. 100 epochs of training were performed for the models analyzed in this study. No consideration of symmetry was given to the cryoDRGN model during training unless otherwise specified. In order to produce a series of volumes for section 2.7, we sampled 50 volumes along the first principal component defined by the VAE latent space of the model from Figure 4.

#### Manifold Embedding

Manifold embedding was run with the following protocol. An aperture size of 4 and an object diameter of 160 Å resulted in sufficiently populated projection directions with no noticeable artifacts from global angular rotations. To avoid the inclusion of noisey projection directions with too few particles, projection directions with fewer than 400 particles were discarded from analysis. No projection directions were discarded for having too many particles. This resulted in 89 PrDs being included in the analysis. The ‘sense’ of the motions was determined by eye. No consideration of symmetry was made during the use of Manifold Embedding unless otherwise specified. In order to produce a series of volumes for section 2.7, we sampled 50 volumes along the conformational coordinate discovered by the manifold embedding calculations in Figure 5.

A new actively maintained version of ManifoldEM can be accessed at the following link https://github.com/flatironinstitute/manifoldem.

### 4.3 Molecular Dynamics and Ensemble Reweighting

#### System Building and MD Simulation Protocols

The source protein structure for this work was the atomic model associated with the raw dataset we have analyzed (PDB ID: 5IRX) [20]. The model for the cytosolic Ankyrin Repeat Domains (PDB ID: 2PNN) was docked into the low resolution electron density of the cryoEM density map (EMDB: 8117). The missing N and C-termini were built using alphafold models and spliced into the model as well. The conformation of the C-terminus was modeled on the conformation found in PDB ID 7LP9, as this structure resolved important interactions between neighboring ARDs. The tails of the four TMD bound palmitoyl oleoyl glycero phosphocholine (POPC) lipids were built manually, in order to ensure their polar contacts with the protein were preserved. Resiniferatoxin parameters were derived from CGENFF [51].

Minimization via the steepest descent algorithm was performed until the force on all atoms was below forces were below 24 kcal/mol/Å. This was followed by relaxation, where a restraint of 10 kcal/mol/*A*^2^ was placed on all heavy atoms and then the magnitude of this restraint is halved every 200 ps, in 15 iterations. Relaxation and production were run with 1 and 2 fs time steps, respectively. Relaxation was followed by 100 ns of equilibration. During relaxation, a Berendsen thermostat and barostat were applied, and for production a Nosé-Hoover and Parrinello-Rahman thermostat and barostat were applied respectively. Production runs were extended up to 2 *μ*s at 310 K. GROMACS 2022 with the CHARMM36m forcefield was used for all molecular dynamics simulations in this publication [52].

#### Principal Component Analysis on Molecular Dynamics Trajectories

Typically, PCA of MD trajectories is dominated by the motion of large disordered loops, because they have a large variance [53]. However, since the mean structure of these disordered domains is not a good estimate for their position in any given frame, these portions of a protein cannot usually be resolved from electron densities. Hence, we applied a standardization transform to the Cartesian coordinates of atoms coordinates before calculating the PCs of their motion. This helps ignore the fluctuations of disordered loops in favor of large collective motions.

In the context of cryoEM, the conventional scheme for applying PCA to molecular dynamics has some deficiencies. Primarily, it assumes that there is no symmetry in the particle of interest. Hence, we developed a simple scheme to apply the notion of symmetry expansion to the trajectory. The trajectory is cloned *N* times, where *N* is the order of the symmetry group of interest (in this case 4 times for C4 symmetry).

In each of these clones, the chain labels are swapped, to simulate the situation that one chain rather than another underwent a large fluctuation.

Dimensionality reduction is a critical part of the analysis of molecular dynamics trajectories. Principal component Analysis is a popular choice but has well-known limitations [53]. One practical difficulty when using PCAs is that large disordered loops typically dominate the analysis, as they produce the most variance. This is a particular problem when comparing eigenvectors to cryoEM, because such loops are not visible. In order to limit the influence of disordered motion in our case, the following procedure of standardization was developed:

The Cartesian coordinates of the atoms in the system are placed in a 3N vector at time *τ*

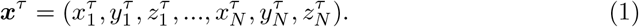

This lets us define a vector of means and variances for each coordinate.

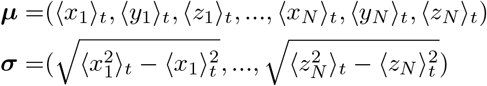

Each 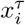 is thus transformed 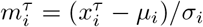. These **m**^*τ*^ are stacked into a matrix to calculate the corresponding singular value decomposition.

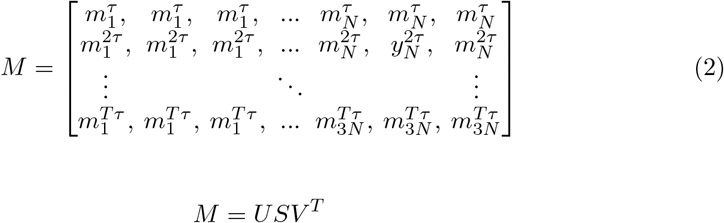

Following this procedure, The columns of V will contain the principal components of M. Code for this procedure can be found in https://github.com/miro-astore/mdanalysis_scripts/pca.py. In order to produce a series of volumes for section 2.7, the first two columns of V were used to derive Cartesian coordinates for 50 atomic models of the backbone of the TRPV1 molecule. These atomic models were then converted into volumes using the *molmap* command in ChimeraX [54].

#### Ensemble Reweighting

Ensemble reweighting leverages experimental data to reweigh a prior conformational ensemble, extracting the conformational probability density suggested by the data. In this study, we employed Bayesian analysis and the cryo-EM particles, in accordance with the mathematical framework detailed in ref. [12], to reweight the 310K ensemble from the MD simulations. We calculated the image-to-structure likelihood using BioEM [55, 56]. We conducted an initial round of BioEM, as described in ref. [57], to determine the best orientations for each image using the reference PDB structure, 36000 quaternions uniformly distributed in SO(3) and variable defocus ∈ [1, 2.6]*μm*. Subsequently, for each image, we generated a list of 8000 quaternions with a finer angular resolution (approximately 2 degrees) around the top 300 orientations from the first round. For the 120 cluster centers obtained from all MD simulations, we employed this refined list of quaternions with BioEM, incorporating the best defocus values from the first round and allowing for variable center translations of 60 pixels along both axes. To prevent misalignment, a low-resolution nanodisc, derived from the reconstructed volume, was added to all atomic structures. The likelihood for each image and each cluster center was calculated within the Ensemble reweighting framework (specifically Eq. ref. [12]) to infer the weight of each cluster center. This inference was performed using Markov chain Monte Carlo (MCMC) with 8 replicas and 20,000 MCMC steps, utilizing the code available at https://github.com/flatironinstitute/Ensemble-reweighting-using-Cryo-EM-particles. The inferred weights were then distributed to the structures from the 310K MD ensemble by nearest neighbor clustering in PC1 and PC2 space.

#### Clustering of MD trajectories

During the reweighting procedure, a subset of structures are chosen from the full ensemble, to limit computational expense. States are typically chosen to be representative of the full ensemble, details can be found in [5]. For this study, structures were chosen using kmedoids clustering on the PC1 and PC2 variables defined in section 2.5. 100 cluster centers were chosen from simulations performed at 310 K, while 10 cluster centers were chosen from simulations at 200 K and 150 K each, for a total of 120 cluster centers.

## 5 Comparing Series of Volumes with CryoSVD

### 5.1 Embedding volumes in a common basis with singular value decomposition

Singular value decomposition (SVD) is a linear algebra technique that, under some conditions, is equivalent to principal component analysis. In short, SVD gives us an orthogonal basis decomposition for the both the rows and the columns of a given matrix. If such rows or columns are mean-removed, then we recover PCA. Here we use SVD to estimate the similarity of the volume series from the different heterogeneity methods by embedding all volumes to a common low dimensional space.

To run our pipeline, we start by flattening and normalizing all the volumes (*X*) so that their l2-norm is 1, 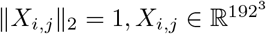 for all *i* = 1, …, *N*_series_ and *j* = 1, …, *N*_*v*_. This will make it so that all the decompositions are on the same scale. Then we remove the mean volume from each series, that is, for every method we define the mean-removed matrix 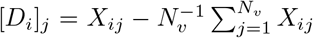, with 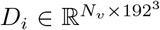. Lastly, we compute an individual economy-SVD for each of these matrices, that is we obtain 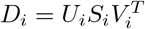, where 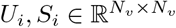, and 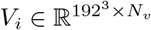.

### 5.2 Comparing the SVD of different volume series

To compare the different basis obtained by SVD we adapt a metric introduced in ref. [13], named the proportion of captured variance (PCV). This metric quantifies the proportion of captured variance between the subspace spanned by the left singular vectors (*V*_*i*_) of two volume series. Let (*U*_*i*_, *S*_*i*_, *V*_*i*_) and (*U*_*j*_, *S*_*j*_, *V*_*j*_) be the SVD from the i-th and j-th volume series respectively. The PCV is defined as

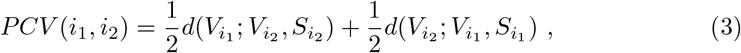

where 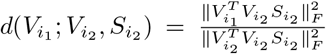 and 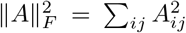 denoting the Frobenius norm. Subspaces that capture the same variance will have a PCV of 1. This metric will allow us to compare different subspaces, while taking into account the level of importance given by the singular values.

### 5.3 Finding a common embedding for a collection of volume series with CryoSVD

When we compute the SVD for a single volume series, we are obtaining the main features ordered based on their variance across the series. Now we are interested in comparing the features across different series. To do this we perform a second round of SVD using the eigenvectors (*V*_*i*_) found previously. From the first round, we have that we can represent any mean-removed volume as

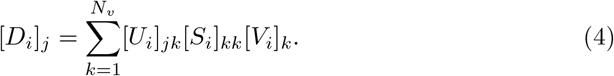

Now, if we perform a second round of SVD by weighting all the *V*_*i*_’s by their respective singular values [*S*_*i*_]_*kk*_ and stacking them, we have yet another basis representation

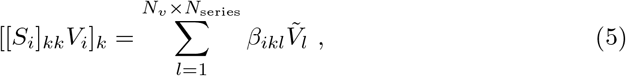

where the vectors 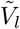 correspond to the left-singular vectors of the second round of SVD. Lastly, we can combine both series to represent the original volumes in the basis defined by 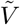, which gives us

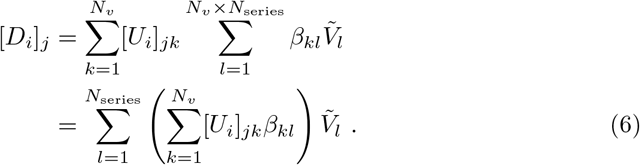

The coefficients 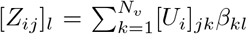, with *l* = 1, …, *N*_*v*_ *× N*_*series*_ are the coordinates of the mean-removed volume [*D*_*i*_]_*j*_ in the lower dimensional space given by the second round of SVD.

### 5.4 Detecting compositional heterogeneity by inspecting power spectrums of volume series

Due to the insensitivity of a Fourier basis to translational degrees of freedom, inspection of the power spectrum for a volume series is a natural way to study the amount of compositional heterogeneity in the series. Series which only capture conformational degrees of freedom will have relatively small changes in their power spectra compared to series with purely compositional heterogeneity. To compare these changes between volume series, we first calculate the radially averaged power spectrum PS_*i*_(*k*) for each volume *i* in a series *i* = 1, …, *N*_*v*_ with *k* the radial reciprocal component and *N*_*v*_ the total number of volumes in the series. Then, we normalize those power spectrums according to the mean of the values at *k* = 0 PS_*i*_(*k*)(*k*) = PS_*i*_, and calculate the variance of the series for each radial frequency 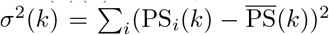 where 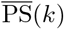 is the mean of the power spectra of the series at radial frequency *k*. In figure over frequencies, i.e., 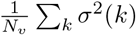, for each volume series, normalised by the number 8D, we present the total power spectrum variance that is the sum of the variances of volumes in the series. By adding up the amount of variance for the series across its power spectrum, we can compare the relative amounts of compositional heterogeneity in different volume series.

The CryoSVD pipeline can be accessed at the following link https://github.com/flatironinstitute/cryoSVD.

## 6 Acknowledgements

The Flatiron Institute is a division of the Simons Foundation. This project is indebted to the support of Joachim Frank, in whose lab related analyses to those presented here were initially performed, and who provided feedback on later stages of the manuscript. We would also like to thank Geoffrey Woollard for his guidance in the development of the cryoSVD pipeline. We would also like to thank the Flatiron Institute’s Scientific Computing Core and Géraud Krawezik in particular for their continuous technical support throughout this project.

## 7 Author Contributions

MA and SMH conceptualized and carried out the project. MA, RB, DS, and PC contributed to the development and application of novel analysis pipelines. MA, PC, and SMH prepared figures. MA and SMH wrote the manuscript, with edits and feedback from all authors.

## 9 Supplementary Material

**Fig. S1.**
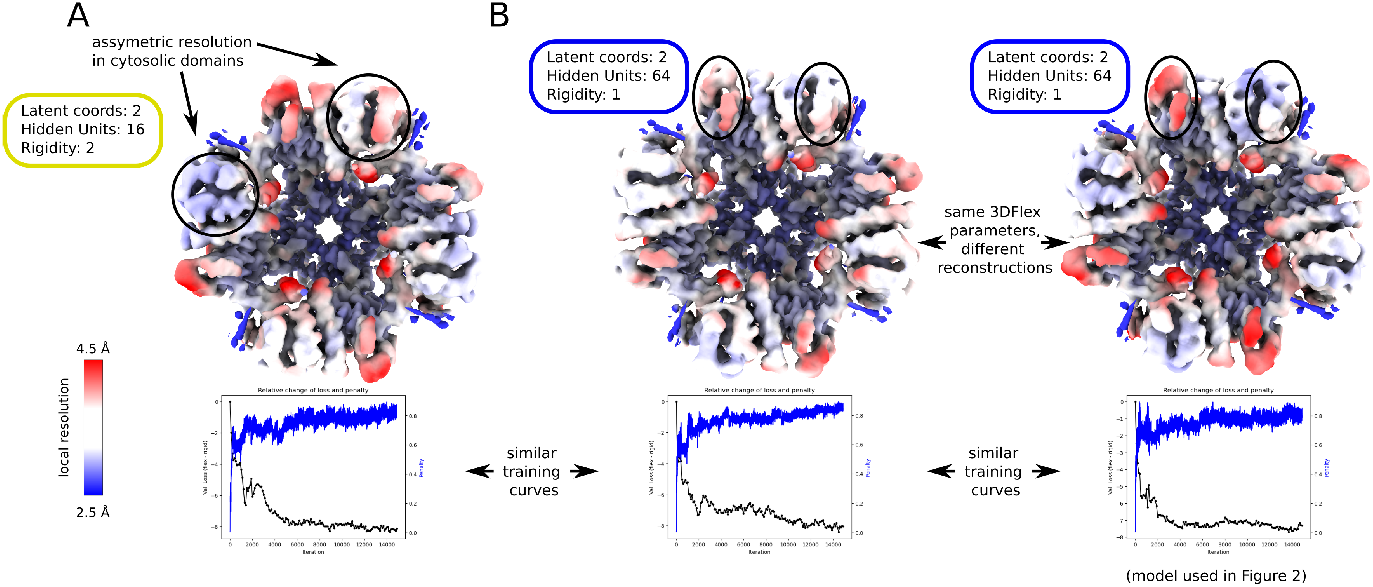
Independently trained 3DFlex models may result in different learned motions and different reconstructions. A) A 3DFlex model trained for a symmetric molecule may result in symmetric units of the molecule having qualitatively different reconstructions with different resolutions. These differences are less pronounced with more flexible models. B) Independent training of different 3DFlex models with the same initial parameters may produce qualitatively different reconstructions from different learned deformations.

**Fig. S2.**
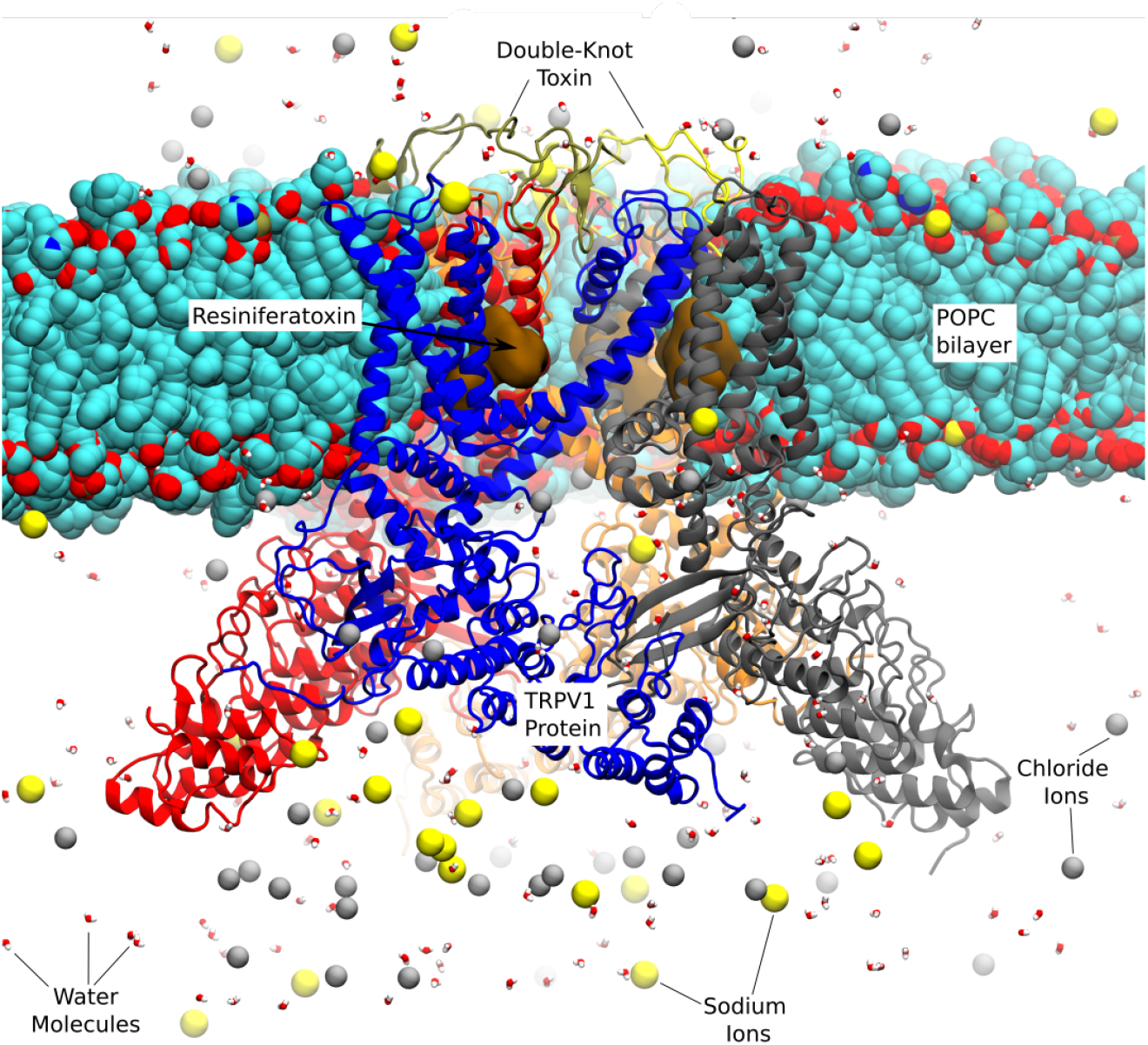
TRPV1 bound to agonist ligands and inserted into a lipid bilayer. The TRPV1 ion channel was inserted into a POPC membrane and solvated with TIP3P water and ionized with sodium and chloride to physiological concentration.

**Fig. S3.**
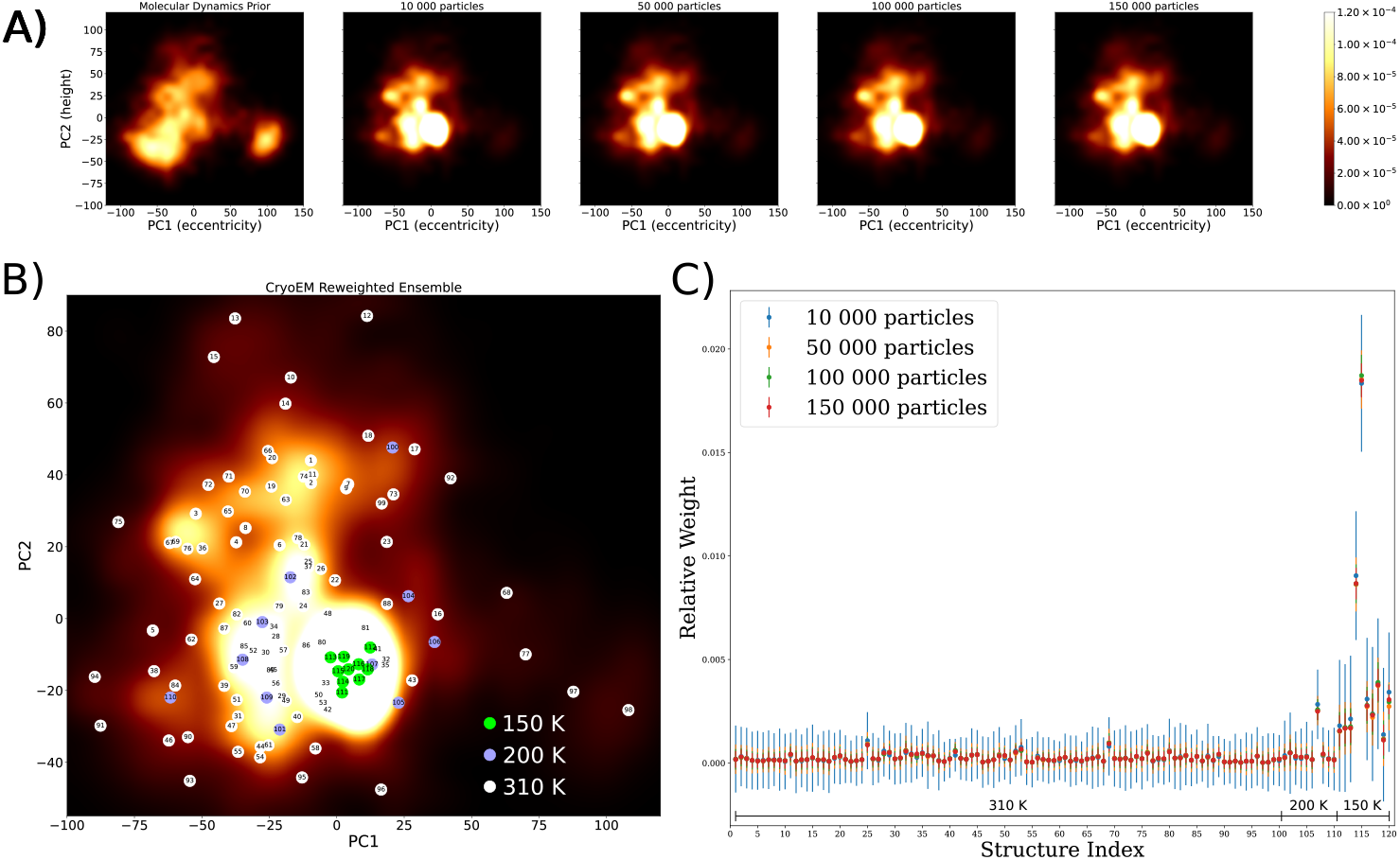
Analysis of convergence of Bayesian ensemble reweighting. A) The prior probability density for molecular dynamics is reweighted in light of varying numbers of particles from the TRPV1 cyroEM dataset. After just 50 000 particles, the estimates for the probability surface have converged. B) The positions of the different structures used for clustering the molecular dynamics trajectories. Most structures cluster around the region occupied by the reference ligand-bound open structure. C) The individual weights estimated by Bayesian ensemble reweighting, with different numbers of particles. This confirms the fast convergence of the algorithm.

**Fig. S4.**
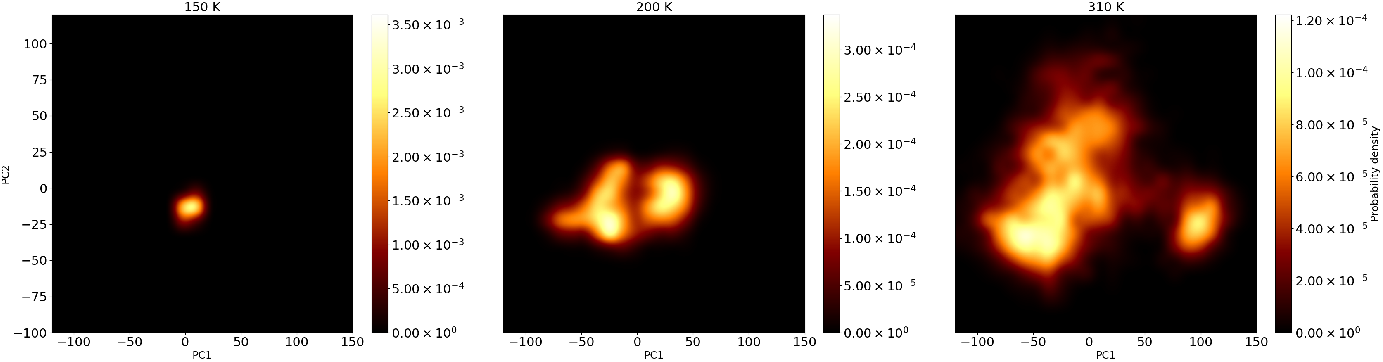
Sampled states of TRPV1 at different temperatures in MD simulation. At lower temperatures, many fewer states are sampled.

**Fig. S5.**
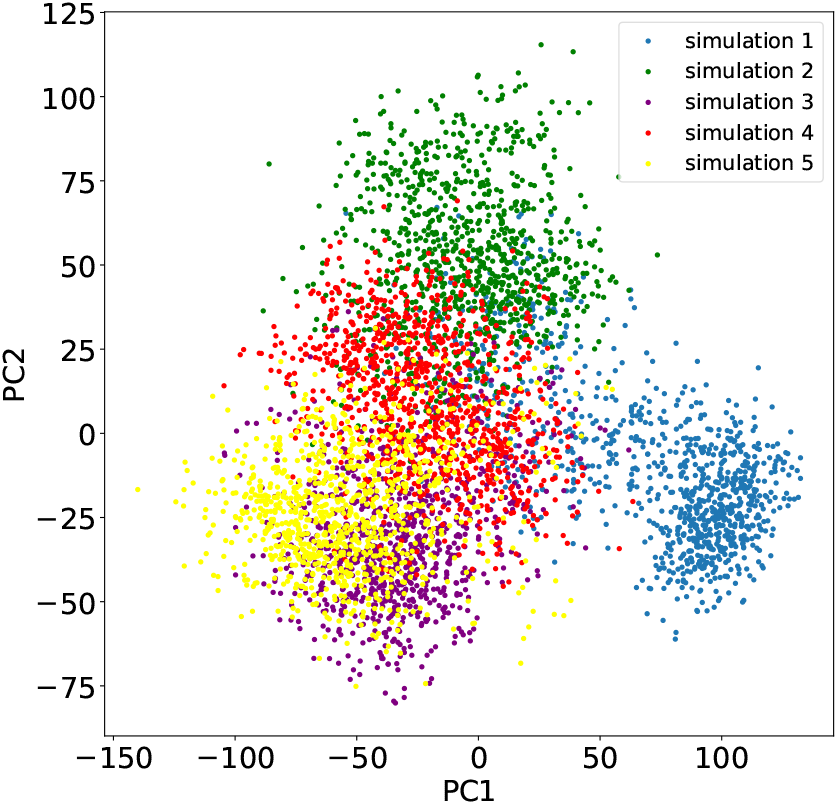
MD replicates of TRPV1 at 310 K. Some replicates sample different parts of the phase space.

**Fig. S6.**
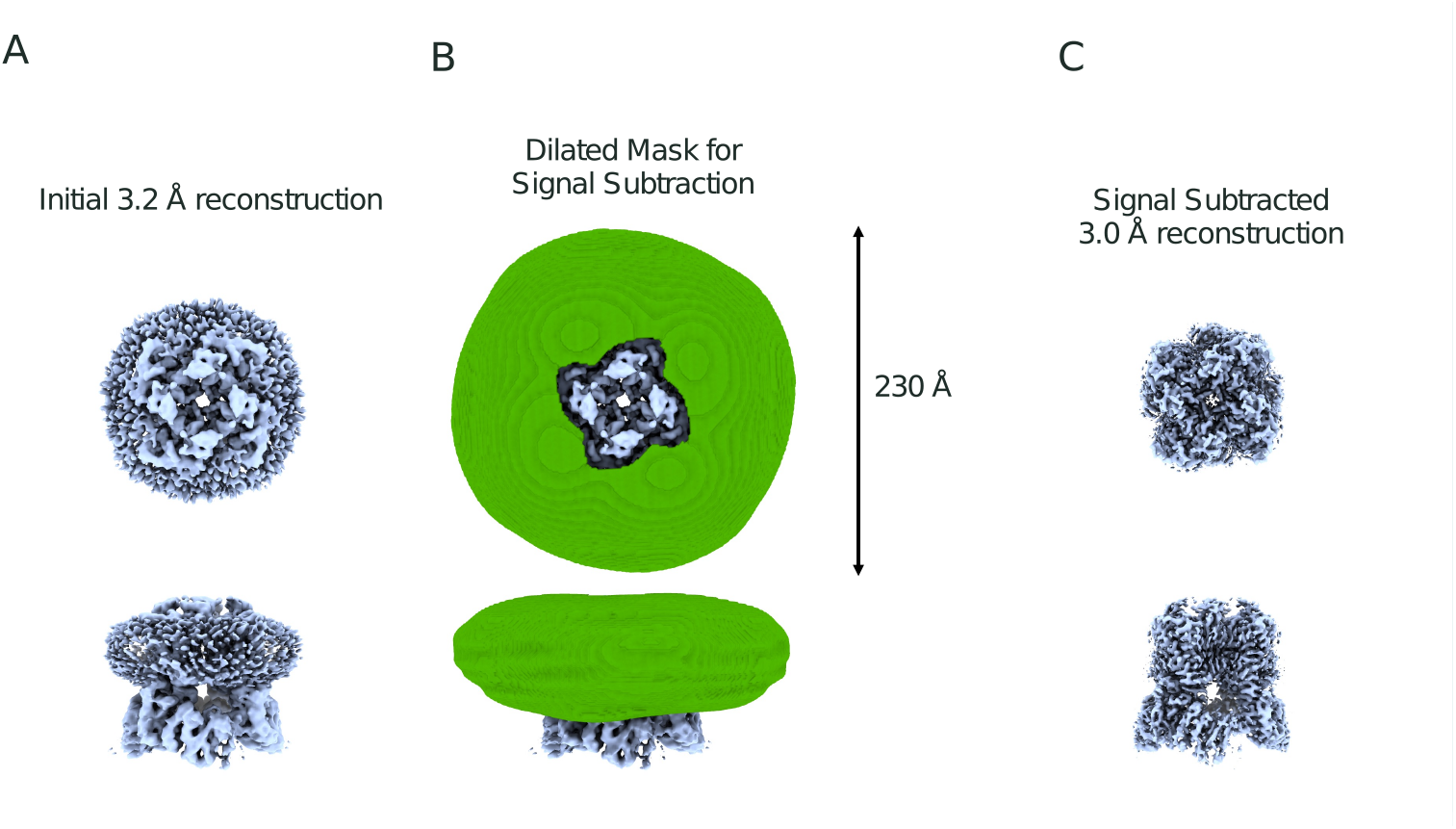
Signal subtraction was performed on the particles stacks before analysis with heterogeneity tools. A) The nanodisc is clearly visible in the homogeneous reconstruction of TRPV1. B) To remove its presence from the results of heterogeneity methods, a dilated nanodisc mask was constructed for signal subtraction. C) This results in the nanodisc being absent from the homogeneous reconstruction.

**Fig. S7.**
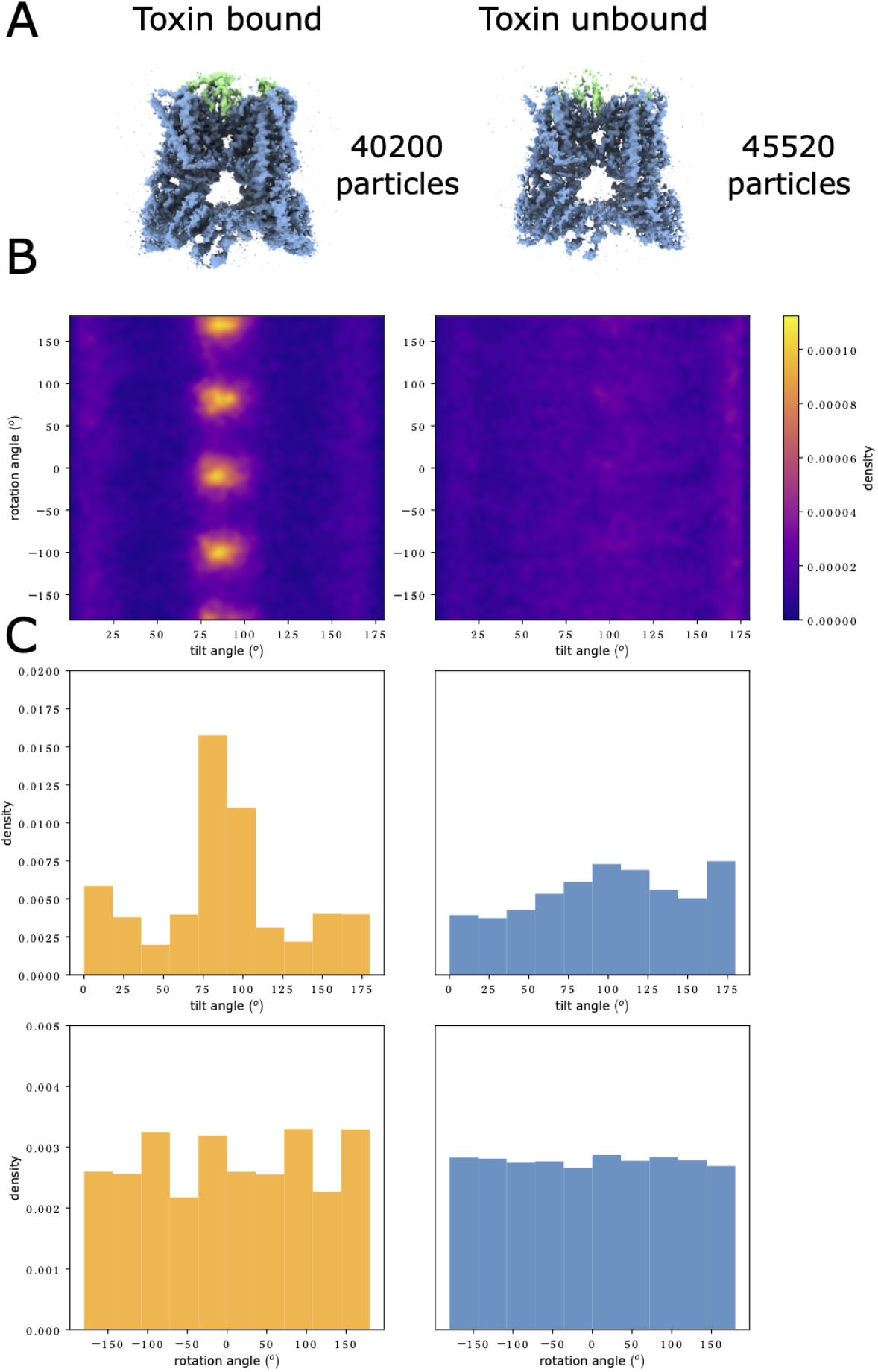
The distribution of orientations for different compositional states of particles. A) Volumes reconstructed from the cryoDRGN latent space with and without high occupancies of DkTX. B) A heat map showing the orientation distribution of particles in both structures. the toxin unbound state has a nearly uniform orientation distribution. C) Histograms of the sum of the two orientation angles along both axes.

**Fig. S8.**
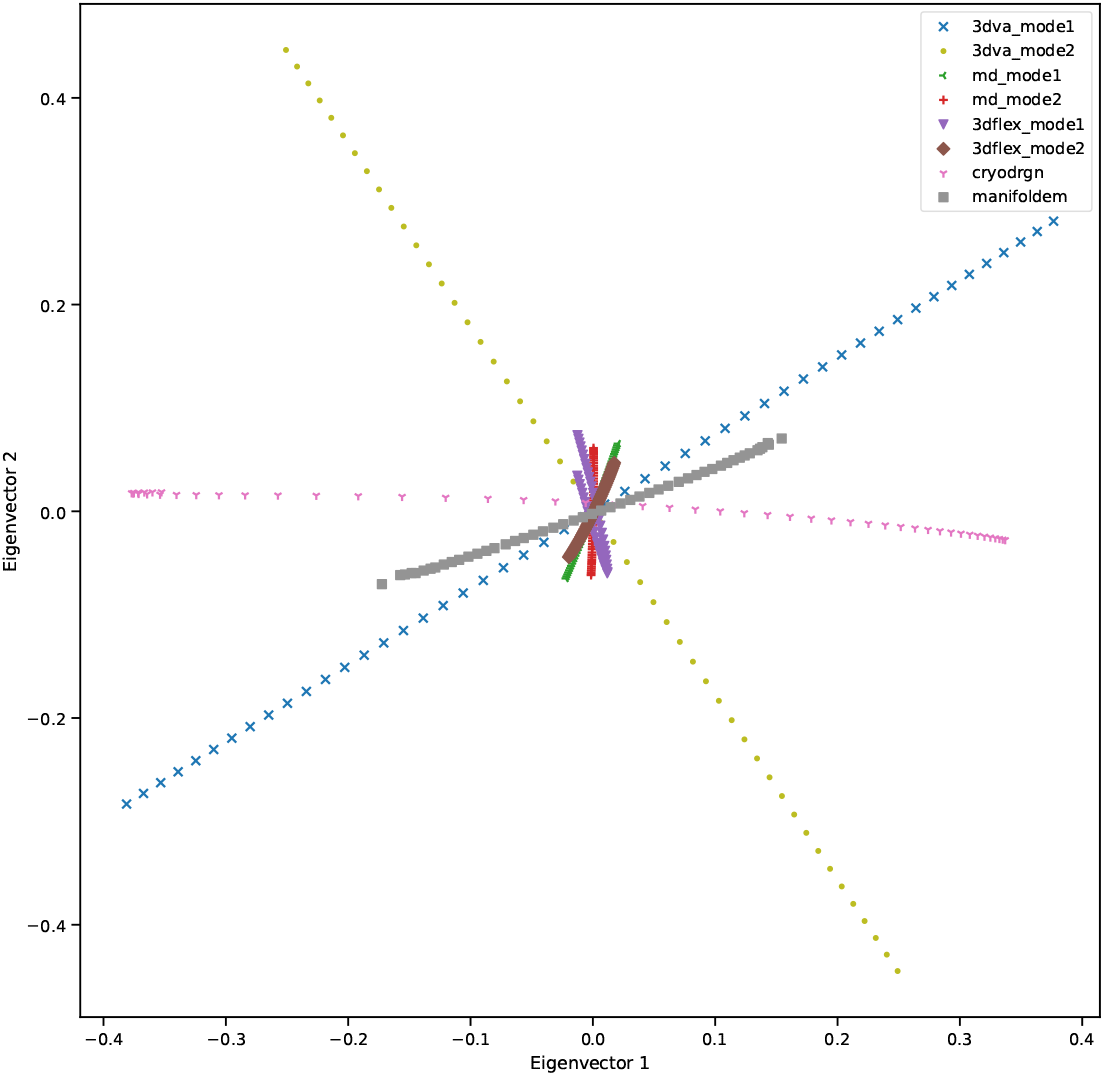
Volume series in a common embedding. The volume series of each method embedded in a common linear basis derived from two rounds of singular value decomposition.

## List of Supplemental Movies

1. “closed_open_morph.mp4” A linear morph between two volumes. The first, (EMDB 8117) represents TRPV1 in the toxin bound open state. The second, (EMD 8118) represents TRPV1 in the apo closed state.
2. “3dflex_trpv1_mode[1,2].mp4” a series of volumes generated from the principal components of the 3DFlex latent space illustrated in figure 2 C.
3. “3dva_mode[1,2]_nanodisc.mp4” a series of volumes generated from the principal components of a 3DVA latent space trained without signal subtraction of the nanodisc.
4. “3dva_mode[1,2].mp4” a series of volumes generated from the first principal component of the 3DVA latent space illustrated in figure 3 C.
5. “cryodrgn_nanodisc_pc1.mp4” a series of volumes generated from the first principal component of a cryoDRGN VAE trained without signal subtraction.
6. “cryodrgn_published_stack_pc[1,2].mp4” a series of volumes generated from the principal components of the latent space of the cryodrgn model depicted in figure 4 B.
7. “cryodrgn_repicked_stack_pc[1,2].mp4” a series of volumes generated from the principal components of the latent space of a cryodrgn model trained on a stack of 525, 910 particles picked from the original micrographs.
8. “cryodrgn_repicked_stack_no_signal_subtraction_pc1.mp4” a series of volumes 930 generated from the principal components of the latent space of a cryodrgn model trained on a stack of 525, 910 particles picked from the original micrographs, without signal subtraction of the nanodisc.
9. “cryodrgn_particles_morph_published_stack.mp4” a linear interpolation between the two reconstructions obtained from the particles indicated by the cryodrgn model to have distinct compositional states.
10. “cryodrgn_particles_morph_repicked_stack.mp4” a linear interpolation between the two reconstructions obtained from the particles indicated by the cryodrgn model trained on 525, 910 particles to have distinct compositional states.
11. “manifold_em_morph_volumes.mp4” a volume series generated along the conformational coordinate of the manifold embedding calculations depicted in figure 5 D.
12. “MD_cryoEM_motions_comparison_trpv1.mp4” A side by side comparison of the eigenvectors from Molecular Dynamics simulations illustrated in figure7 and the volume series along the principal components calculated by 3D Variability analysis in figure 3A-B.
13. “supplementary_3dflex_independent_trainings_mode_[1,2].mp4” volumes series along the principal components of models with identical hyper parameters and training data as the model trained in figure 2. This model has learned slightly different deformations to the previous model.
14. “3dva 3dclass particles mode [1,2].mp4” volumes series generated from the principal components of 3D Variability Analysis calculations performed on a stack of 57 602 high quality particles from 3d classification.
15. “manifoldem_signal_subtraction_effects.mp4” The effects of masking and signal subtraction on the NLSA eigenvectors of Manifold Embedding analysis.

## References

[1] Bank, R.P.D.: PDB Statistics: Number of Released PDB Structures per Year. https://www.rcsb.org/stats/all-released-structures Accessed 2024-09-06

[2] Callaway, E.: The revolution will not be crystallized: a new method sweeps through structural biology. Nature 525(7568), 172–174 (2015) 10.1038/525172a. Publisher: Nature Publishing Group. Accessed 2024-09-06

[3] Heel, M., Frank, J.: Use of multivariates statistics in analysing the images of biological macromolecules. Ultramicroscopy 6(1), 187–194 (1981) 10.1016/S0304-3991(81)80197-0. Accessed 2024-04-17

[4] Li, Y., Zhang, R., Wang, C., Forouhar, F., Clarke, O.B., Vorobiev, S., Singh, S., Montelione, G.T., Szyperski, T., Xu, Y., Hunt, J.F.: Oligomeric interactions maintain active-site structure in a noncooperative enzyme family. The EMBO Journal 41(17), 108368 (2022) 10.15252/embj.2021108368. Publisher: John Wiley & Sons, Ltd. Accessed 2024-04-17

[5] Tang, W.S., Zhong, E.D., Hanson, S.M., Thiede, E.H., Cossio, P.: Conformational heterogeneity and probability distributions from single-particle cryo-electron microscopy. Current Opinion in Structural Biology 81, 102626 (2023) 10.1016/j.sbi.2023.102626. Accessed 2023-12-14

[6] Dashti, A., Schwander, P., Langlois, R., Fung, R., Li, W., Hosseinizadeh, A., Liao, H.Y., Pallesen, J., Sharma, G., Stupina, V.A., Simon, A.E., Dinman, J.D., Frank, J., Ourmazd, A.: Trajectories of the ribosome as a Brownian nanoma-chine. Proceedings of the National Academy of Sciences 111(49), 17492–17497 (2014) 10.1073/pnas.1419276111. Publisher: Proceedings of the National Academy of Sciences. Accessed 2023-08-25

[7] Zhong, E.D., Bepler, T., Berger, B., Davis, J.H.: CryoDRGN: reconstruction of heterogeneous cryo-EM structures using neural networks. Nature Methods 18(2), 176–185 (2021) 10.1038/s41592-020-01049-4. Number:2 Publisher: Nature Publishing Group. Accessed 2022-12-22

[8] Schwab, J., Kimanius, D., Burt, A., Dendooven, T., Scheres, S.H.W.: DynaMight: estimating molecular motions with improved reconstruction from cryo-EM images. bioRxiv. Pages: 2023.10.18.562877 Section: New Results (2023). 10.1101/2023.10.18.562877. https://www.biorxiv.org/content/10.1101/2023.10.18.562877v1 Accessed 2024-08-25

[9] Chen, M., Ludtke, S.J.: Deep learning-based mixed-dimensional Gaussian mixture model for characterizing variability in cryo-EM. Nature Methods 18(8), 930–936 (2021) 10.1038/s41592-021-01220-5. Publisher: Nature Publishing Group. Accessed 2024-09-06

[10] Punjani, A., Fleet, D.J.: 3DFlex: determining structure and motion of flexible proteins from cryo-EM. Nature Methods 20(6), 860–870 (2023) 10.1038/s41592-023-01853-8. Number: 6 Publisher: Nature Publishing Group. Accessed 2023-08-25

[11] Herreros, D., Lederman, R.R., Krieger, J.M., Jiménez-Moreno, A., Martínez, M., Myška, D., Strelak, D., Filipovic, J., Sorzano, C.O.S., Carazo, J.M.: Estimating conformational landscapes from Cryo-EM particles by 3D Zernike polynomials. Nature Communications 14(1), 154 (2023) 10.1038/s41467-023-35791-y. Publisher: Nature Publishing Group. Accessed 2024-09-06

[12] Tang, W.S., Silva-Sánchez, D., Giraldo-Barreto, J., Carpenter, B., Hanson, S.M., Barnett, A.H., Thiede, E.H., Cossio, P.: Ensemble Reweighting Using CryoEM Particle Images. The Journal of Physical Chemistry B 127(24), 5410–5421 (2023) 10.1021/acs.jpcb.3c01087. Publisher: American Chemical Society. Accessed 2023-08-26

[13] Gilles, M.A.T., Singer, A.: Cryo-EM Heterogeneity Analysis using Regularized Covariance Estimation and Kernel Regression (2023). 10.1101/2023.10.28.564422. http://biorxiv.org/lookup/doi/10.1101/2023.10.28.564422 Accessed 2024-08-14

[14] Punjani, A., Fleet, D.J.: 3D variability analysis: Resolving continuous flexibility and discrete heterogeneity from single particle cryo-EM. Journal of Structural Biology 213(2), 107702 (2021) 10.1016/j.jsb.2021.107702. Accessed 2023-08-25

[15] Cao, E., Liao, M., Cheng, Y., Julius, D.: TRPV1 structures in distinct conformations reveal mechanisms of activation. Nature 504(7478), 113–118 (2013) 10.1038/nature12823. Accessed 2024-04-26

[16] Caterina, M.J., Schumacher, M.A., Tominaga, M., Rosen, T.A., Levine, J.D., Julius, D.: The capsaicin receptor: a heat-activated ion channel in the pain pathway. Nature 389(6653), 816–824 (1997) 10.1038/39807. Accessed 2024-08-05

[17] Clapham, D.E., Miller, C.: A thermodynamic framework for understanding temperature sensing by transient receptor potential (TRP) channels. Proceedings of the National Academy of Sciences 108(49), 19492–19497 (2011) 10.1073/pnas.1117485108. Publisher: Proceedings of the National Academy of Sciences. Accessed 2024-04-01

[18] Yeh, F., Jara-Oseguera, A., Aldrich, R.W.: Implications of a temperature-dependent heat capacity for temperature-gated ion channels. Proceedings of the National Academy of Sciences of the United States of America 120(24), 2301528120 (2023) 10.1073/pnas.2301528120

[19] Singh, A.K., McGoldrick, L.L., Demirkhanyan, L., Leslie, M., Zakharian, E., Sobolevsky, A.I.: Structural basis of temperature sensation by the TRP channel TRPV3. Nature structural & molecular biology 26(11), 994–998 (2019)10.1038/s41594-019-0318-7. Accessed 2024-05-07

[20] Gao, Y., Cao, E., Julius, D., Cheng, Y.: TRPV1 structures in nanodiscs reveal mechanisms of ligand and lipid action. Nature 534(7607), 347–351 (2016) 10.1038/nature17964

[21] Zhang, K., Julius, D., Cheng, Y.: Structural snapshots of TRPV1 reveal mechanism of polymodal functionality. Cell 184(20), 5138–515012 (2021) 10.1016/j.cell.2021.08.012

[22] Sorzano, C.O.S., Carazo, J.M.: Principal component analysis is limited to lowresolution analysis in cryoEM. Acta Crystallographica Section D: Structural Biology 77(6), 835–839 (2021) 10.1107/S2059798321002291. Publisher: International Union of Crystallography. Accessed 2024-04-02

[23] Schwander, P., Fung, R., Ourmazd, A.: Conformations of macromolecules and their complexes from heterogeneous datasets. Philosophical Transactions of the Royal Society B: Biological Sciences 369(1647), 20130567 (2014) 10.1098/rstb.2013.0567. Accessed 2023-02-23

[24] Sztain, T., Ahn, S.-H., Bogetti, A.T., Casalino, L., Goldsmith, J.A., Seitz, E., McCool, R.S., Kearns, F.L., Acosta-Reyes, F., Maji, S., Mashayekhi, G., McCammon, J.A., Ourmazd, A., Frank, J., McLellan, J.S., Chong, L.T., Amaro, R.E.: A glycan gate controls opening of the SARS-CoV-2 spike protein. Nature Chemistry 13(10), 963–968 (2021) 10.1038/s41557-021-00758-3. Publisher: Nature Publishing Group. Accessed 2024-07-11

[25] Mashayekhi, G., Vant, J., Polavarapu, A., Ourmazd, A., Singharoy, A.: Energy landscape of the SARS-CoV-2 reveals extensive conformational heterogeneity. Current Research in Structural Biology 4, 68–77 (2022) 10.1016/j.crstbi.2022.02.001. Accessed 2024-07-11

[26] Dashti, A., Mashayekhi, G., Shekhar, M., Ben Hail, D., Salah, S., Schwander, P., Georges, A., Singharoy, A., Frank, J., Ourmazd, A.: Retrieving functional pathways of biomolecules from single-particle snapshots. Nature Communications 11(1), 4734 (2020) 10.1038/s41467-020-18403-x. Number: 1 Publisher: Nature Publishing Group. Accessed 2023-08-25

[27] Seitz, E., Acosta-Reyes, F., Maji, S., Schwander, P., Frank, J.: Recovery of Conformational Continuum From Single-Particle Cryo-EM Images: Optimization of ManifoldEM Informed by Ground Truth. IEEE Transactions on Computational Imaging 8, 462–478 (2022) 10.1109/TCI.2022.3174801. Conference Name: IEEE Transactions on Computational Imaging

[28] Maji, S., Liao, H., Dashti, A., Mashayekhi, G., Ourmazd, A., Frank, J.: Propagation of Conformational Coordinates Across Angular Space in Mapping the Continuum of States from Cryo-EM Data by Manifold Embedding. Journal of Chemical Information and Modeling 60(5), 2484–2491 (2020) 10.1021/acs.jcim.9b01115. Accessed 2024-07-11

[29] Giraldo-Barreto, J., Ortiz, S., Thiede, E.H., Palacio-Rodriguez, K., Carpenter, B., Barnett, A.H., Cossio, P.: A Bayesian approach to extracting free-energy profiles from cryo-electron microscopy experiments. Scientific Reports 11(1), 13657 (2021) 10.1038/s41598-021-92621-1. Publisher: Nature Publishing Group. Accessed 2024-07-11

[30] Forsberg, B.O., Shah, P.N.M., Burt, A.: A robust normalized local filter to estimate compositional heterogeneity directly from cryo-EM maps. Nature Communications 14(1), 5802 (2023) 10.1038/s41467-023-41478-1. Publisher: Nature Publishing Group. Accessed 2024-03-10

[31] Jeon, M., Raghu, R., Astore, M., Woollard, G., Feathers, R., Kaz, A., Hanson, S.M., Cossio, P., Zhong, E.D.: CryoBench: Diverse and challenging datasets for the heterogeneity problem in cryo-EM. arXiv. 2408.05526 [cs, q-bio] (2024). 10.48550/arXiv.2408.05526. http://arxiv.org/abs/2408.05526 Accessed 2024-08-20

[32] Miro Astore, Geoffrey Woollard, David Silva, Khanh Dao Duc, Nikolaus Grigorieff, Pilar Cossio, Sonya M Hanson: The Inaugural Flatiron Institute Cryo-EM Heterogeneity Community Challenge (2024) 10.17605/OSF.IO/8H6FZ. Publisher: OSF. Accessed 2024-08-20

[33] Zhu, J., Zhang, Q., Zhang, H., Shi, Z., Hu, M., Bao, C.: A minority of final stacks yields superior amplitude in single-particle cryo-EM. Nature Communications 14(1), 7822 (2023) 10.1038/s41467-023-43555-x. Publisher: Nature Publishing Group. Accessed 2024-03-22

[34] Laursen, W.J., Schneider, E.R., Merriman, D.K., Bagriantsev, S.N., Gracheva, E.O.: Low-cost functional plasticity of TRPV1 supports heat tolerance in squirrels and camels. Proceedings of the National Academy of Sciences 113(40), 11342– 11347 (2016) 10.1073/pnas.1604269113. Publisher: Proceedings of the National Academy of Sciences. Accessed 2024-08-20

[35] Gracheva, E.O., Cordero-Morales, J.F., González-Carcacía, J.A., Ingolia, N.T., Manno, C., Aranguren, C.I., Weissman, J.S., Julius, D.: Ganglion-specific splicing of TRPV1 underlies infrared sensation in vampire bats. Nature 476(7358), 88– 91 (2011) 10.1038/nature10245. Publisher: Nature Publishing Group. Accessed 2024-08-20

[36] Ladrón-de-Guevara, E., Dominguez, L., Rangel-Yescas, G.E., Fernández-Velasco, D.A., Torres-Larios, A., Rosenbaum, T., Islas, L.D.: The Contribution of the Ankyrin Repeat Domain of TRPV1 as a Thermal Module. Biophysical Journal 118(4), 836–845 (2020) 10.1016/j.bpj.2019.10.041. Publisher: Elsevier. Accessed 2024-08-23

[37] Tie Li, J.-W.L.: Oscillation of S5 helix under different temperatures in determination of the open probability of TRPV1 channel. Chinese Physics B 29(9), 98701–098701 (2020) 10.1088/1674-1056/aba600. Accessed 2024-08-23

[38] Wen, H., Qin, F., Zheng, W.: Toward elucidating the heat activation mechanism of the TRPV1 channel gating by molecular dynamics simulation. Proteins: Structure, Function, and Bioinformatics 84(12), 1938–1949 (2016) 10.1002/prot.25177. eprint: https://onlinelibrary.wiley.com/doi/pdf/10.1002/prot.25177. Accessed 2024-08-23

[39] Astore, M.A., Pradhan, A.S., Thiede, E.H., Hanson, S.M.: Protein dynamics underlying allosteric regulation. Current Opinion in Structural Biology 84, 102768 (2024) 10.1016/j.sbi.2023.102768. Accessed 2024-08-20

[40] Cooper, A., Dryden, D.T.: Allostery without conformational change. A plausible model. European biophysics journal: EBJ 11(2), 103–109 (1984) 10.1007/BF00276625

[41] Bock, L.V., Grubmüller, H.: Effects of cryo-EM cooling on structural ensembles. Nature Communications 13(1), 1709 (2022) 10.1038/s41467-022-29332-2. Number: 1 Publisher: Nature Publishing Group. Accessed 2023-08-29

[42] Mowry, N.J., Kruger, C.R., Drabbels, M., Lorenz, U.J.: Direct Measurement of the Critical Cooling Rate for the Vitrification of Water (2024). https://arxiv.org/abs/2407.01087v1 Accessed 2024-09-06

[43] Heel, M., Schatz, M.: Fourier shell correlation threshold criteria. Journal of Structural Biology 151(3), 250–262 (2005) 10.1016/j.jsb.2005.05.009. Accessed 2024-09-03

[44] Pb, R., R, H.: Optimal determination of particle orientation, absolute hand, and contrast loss in single-particle electron cryomicroscopy. Journal of molecular biology 333(4) (2003) 10.1016/j.jmb.2003.07.013. Publisher: J Mol Biol. Accessed 2024-09-03

[45] Torino, S., Dhurandhar, M., Stroobants, A., Claessens, R., Efremov, R.G.: Time-resolved cryo-EM using a combination of droplet microfluidics with ondemand jetting. Nature Methods 20(9), 1400–1408 (2023) 10.1038/s41592-023-01967-z. Publisher: Nature Publishing Group. Accessed 2024-08-27

[46] Papasergi-Scott, M.M., Pérez-Hernández, G., Batebi, H., Gao, Y., Eskici, G., Seven, A.B., Panova, O., Hilger, D., Casiraghi, M., He, F., Maul, L., Gmeiner, P., Kobilka, B.K., Hildebrand, P.W., Skiniotis, G.: Time-resolved cryo-EM of G-protein activation by a GPCR. Nature 629(8014), 1182–1191 (2024) 10.1038/s41586-024-07153-1. Publisher: Nature Publishing Group. Accessed 2024-08-27

[47] Zhang, K., Lucas, B., Grigorieff, N.: Exploring the Limits of 2D Template Matching for Detecting Targets in Cellular Cryo-EM Images. Microscopy and Microanalysis 29(Supplement 1), 931 (2023) 10.1093/micmic/ozad067.462. Accessed 2024-08-27

[48] Lucas, B.A., Himes, B.A., Grigorieff, N.: Baited reconstruction with 2D template matching for high-resolution structure determination in vitro and in vivo without template bias. eLife 12 (2023) 10.7554/eLife.90486.1. Publisher: eLife Sciences Publications Limited. Accessed 2024-08-27

[49] Rangan, R., Feathers, R., Khavnekar, S., Lerer, A., Johnston, J.D., Kelley, R., Obr, M., Kotecha, A., Zhong, E.D.: CryoDRGN-ET: deep reconstructing generative networks for visualizing dynamic biomolecules inside cells. Nature Methods 21(8), 1537–1545 (2024) 10.1038/s41592-024-02340-4. Publisher: Nature Publishing Group. Accessed 2024-09-03

[50] Dalal, V., Arcario, M.J., Petroff, J.T., Tan, B.K., Dietzen, N.M., Rau, M.J., Fitzpatrick, J.A.J., Brannigan, G., Cheng, W.W.L.: Lipid nanodisc scaffold and size alter the structure of a pentameric ligand-gated ion channel. Nature Communications 15(1), 25 (2024) 10.1038/s41467-023-44366-w. Publisher: Nature Publishing Group. Accessed 2024-07-11

[51] Vanommeslaeghe, K., Hatcher, E., Acharya, C., Kundu, S., Zhong, S., Shim, J., Darian, E., Guvench, O., Lopes, P., Vorobyov, I., Mackerell, A.D.: CHARMM general force field: A force field for drug-like molecules compatible with the CHARMM all-atom additive biological force fields. Journal of Computational Chemistry 31(4), 671–690 (2010) 10.1002/jcc.21367

[52] Huang, J., Rauscher, S., Nawrocki, G., Ran, T., Feig, M., Groot, B.L., Grubmüller, H., MacKerell Jr, A.D.: CHARMM36m: an improved force field for folded and intrinsically disordered proteins. Nature Methods 14(1), 71–73 (2016) 10.1038/nmeth.4067. Publisher: Nature Publishing Group

[53] Skjaerven, L., Martinez, A., Reuter, N.: Principal component and normal mode analysis of proteins; a quantitative comparison using the GroEL subunit. Proteins: Structure, Function, and Bioinformatics 79(1), 232–243 (2011) 10.1002/prot.22875. eprint: https://onlinelibrary.wiley.com/doi/pdf/10.1002/prot.22875. Accessed 2024-04-09

[54] Meng, E.C., Goddard, T.D., Pettersen, E.F., Couch, G.S., Pearson, Z.J., Morris, J.H., Ferrin, T.E.: UCSF ChimeraX: Tools for structure building and analysis. Protein Science 32(11), 4792 (2023) 10.1002/pro.4792.eprint: https://onlinelibrary.wiley.com/doi/pdf/10.1002/pro.4792. Accessed 2024-09-07

[55] Cossio, P., Hummer, G.: Bayesian analysis of individual electron microscopy images: towards structures of dynamic and heterogeneous biomolecular assemblies. Journal of Structural Biology 184(3), 427–437 (2013) 10.1016/j.jsb.2013.10.006

[56] Cossio, P., Rohr, D., Baruffa, F., Rampp, M., Lindenstruth, V., Hummer, G.: BioEM: GPU-accelerated computing of Bayesian inference of electron microscopy images. Computer Physics Communications 210, 163–171 (2017) 10.1016/j.cpc.2016.09.014. Accessed 2023-08-28

[57] Cossio, P., Allegretti, M., Mayer, F., Müller, V., Vonck, J., Hummer, G.: Bayesian inference of rotor ring stoichiometry from electron microscopy images of archaeal ATP synthase. Microscopy 67(5), 266–273 (2018) 10.1093/jmicro/dfy033. Accessed 2024-07-11

